# The WAC-downWAC domain in the yeast ISW2 nucleosome remodeling complex forms a structural module essential for ISW2 function but not cell viability

**DOI:** 10.1101/2025.04.27.650761

**Authors:** Ashish Kumar Singh, Sabine Ines Grünert, Lena Pfaller, Felix Mueller-Planitz

## Abstract

**Background:** ATP-dependent nucleosome remodeling complexes of the imitation switch (ISWI) family slide and space nucleosomes. The ISWI ATPase subunit forms obligate complexes with accessory subunits whose mechanistic roles remain poorly understood. In baker’s yeast, the Isw2 ATPase subunit associates with Itc1, the orthologue of human ACF1/BAZ1A. Prior data suggested that the genomic deletion of the 374 N-terminal amino acids from Itc1 (hereafter called *itc1*^ΔN^) leads to a gain-of-toxic-function phenotype with severe growth defects in the BY4741 genetic background, possibly due to defective nucleosome spacing activity of the mutant complex.

**Results:** Here we show that the deletion encompasses a novel motif termed downWAC that forms a conserved structural module with the N-terminal WAC domain. The module is predicted to interact with DNA. However, it does not form a stable interaction interface with the remainder of the complex. Instead, it is connected through a long disordered polypeptide linker to the remainder of the complex. Curiously, the *itc1*^ΔN^ allele does not lead to measurable growth defects in haploid BY4741 and diploid BY4743 strains. It also does not alter genome-wide nucleosome organization in wild-type cells. To rule out that potentially redundant remodeling factors obscure *itc1*^ΔN^-associated phenotypes, we repeated experiments in cells devoid of *ISW1* and *CHD1* remodelers with the same results. Only at known target genes of the ISW2 complex was the nucleosome organization perturbed in *itc1*^ΔN^ cells.

**Conclusions:** We conclude that the deletion of Itc1 N-terminus is indistinguishable from the full deletion of either *ITC1* or *ISW2*. As such, *itc1*^ΔN^ should be considered a null allele of *ISW2*. We propose a model, in which the WAC-downWAC module, along with a flexible protein linker, helps ISW2 in searching for its target genes and positioning +1-nucleosomes.

## Background

Nucleosome positions in eukaryotes are precisely controlled by numerous highly conserved nucleosome remodelling enzymes. Most of these enzymes can reposition nucleosomes along DNA in a process termed nucleosome sliding [1–6]. Some, including the ISWI and CHD1 classes of remodelers, can also equilibrate the distances between nucleosomes *in vivo* and *in vitro* [7–11], a process that is called nucleosome spacing. Accurate nucleosome positioning is crucial for regulating gene expression and maintaining chromatin structure [12].

Remodeling enzymes tend to form multi-subunit complexes. The *Saccharomyces cerevisiae* ISW2 remodeling complex, for instance, consists of the ISWI-type ATPase subunit Isw2 and a large accessory subunit called Itc1 (Figs. 1A, B) [13,14]. Two histone-fold containing proteins, Dls1 and Dpb4, may also associate with the ISW2 complex [15,16]. ISW2 slides and spaces nucleosomes *in vitro* and *in vivo* [10,13,17–19].

**Fig. 1.**
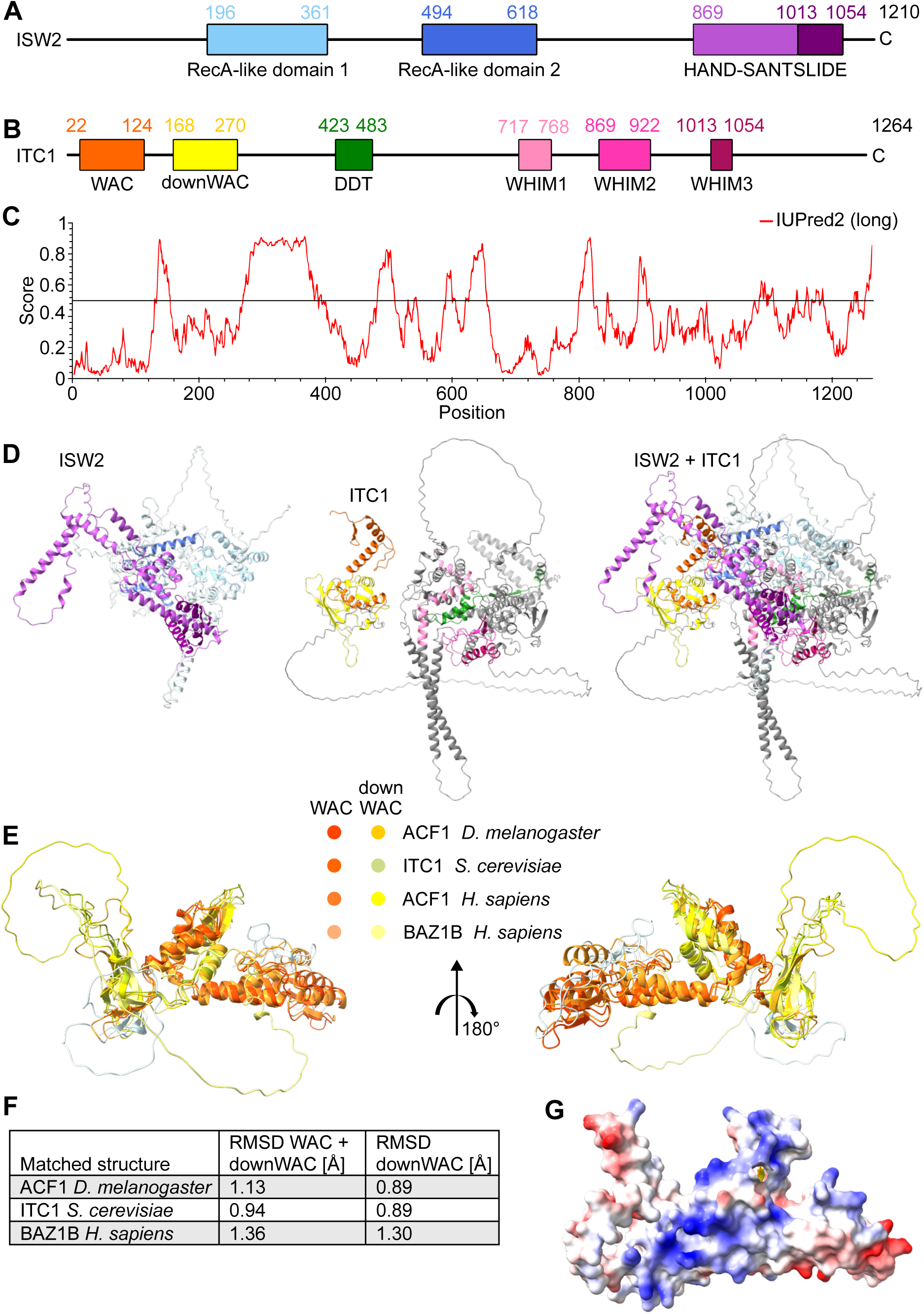
The downWAC motif in the N-terminal regions of *Sc*Itc1 and *Hs*ACF1 forms a structurally conserved module with the adjacent WAC domain. **(A)** Domain organization of *Sc*Isw2. (B) Domain organization of *Sc*Itc1. (C) Prediction of protein disorder using IUPred2A of *Sc*Itc1. (D) Structural model of the *Sc*ISW2 complex generated by AlphaFold 3. The left, middle and right panels show *Sc*Isw2, *Sc*Itc1, or both, respectively. (E) Superimposed AlphaFold 3 predictions of the WAC-downWAC modules across multiple homologous proteins. (F) Structural similarities of the WAC-downWAC modules shown in the previous panel. Shown are root-mean-square deviations (RMSD) to *Hs*Acf1. (G) Electrostatic surface potential of the WAC-downWAC module in *Sc*Itc1. Note the positive surface (blue), which could help in DNA binding, as suggested before [26,27].

The ISW2 complex is orthologous to the human and *Drosophila melanogaster* (*Dm*) ACF/CHRAC complexes. These complexes contain the accessory Acf1 subunit (also known in humans as BAZ1A), which possesses homology to yeast Itc1. Like yeast ISW2, ACF/CHRAC complexes slide and space nucleosomes *in vitro* and *in vivo* [20–25]. The role of the accessory subunits of these complexes, Itc1 in *Saccharomyces cerevisiae* (*Sc*) and Acf1 in metazoans, remain mostly unclear.

Previous work pointed to a critical functional role of the N-terminal region of Itc1 and Acf1. This region includes a WAC (WSTF/Acf1/Cbp146) motif (Fig. 1B). Deletion of the WAC motif in *Drosophila* Acf1 lowers DNA affinity and modestly slows down activity of the complex [26]. Strikingly, deletion of the 371 N-terminal amino acids of *Homo sapiens* (*Hs*) Acf1 led to severe catalytic defects *in vitro*. This deletion quantitatively abolished the ability of the *Hs*ACF complex to measure the length of DNA that flanks the nucleosome, the so called linker DNA *in vitro* [27]. Without sensing the length of linker DNA, *Hs*ACF should no longer be able to space nucleosomes *in vivo*.

Experiments with the orthologous yeast complex supported a critical functional role of the N-terminal region of *Hs*Acf1. The analogous deletion in *Sc*Itc1 (amino acids 2–374; hereafter called *itc1*^ΔN^) (Fig. 1B) dramatically reduced cellular growth in the BY4741 yeast background [27]. The strong growth phenotype of *itc1*^ΔN^ was curious given that neither *ITC1* nor its binding partner *ISW2* are essential genes, suggesting that *itc1*^ΔN^ causes a gain of toxic function in the BY4741 yeast background. Surprisingly, another study did not detect lethality induced by the N-terminal deletion of Itc1 in the W303 yeast strain [28]. Perhaps deletion of Itc1’s N-terminus is toxic in one but not the other genetic background. Different reliance of yeast strains on the presence of remodelers is not unprecedented: whereas the INO80 remodeler is dispensable in the BY4741 background, it is essential in the W303 background [29].

Here we sensitively test the hypothesis that the absence of Itc1’s N-terminus leads to a gain of toxic function, possibly through the mutant remodeler producing irregularly spaced nucleosomes. Unexpectedly, our results demonstrate that the deletion of the N-terminus of Itc1 is well tolerated in the BY4741 yeast background with no measurable growth defect. We ruled out that other remodelers that space nucleosomes interfered with our results. We measured the genome-wide nucleosome organization and did not find evidence for *itc1*^ΔN^ causing abnormal levels of irregular nucleosome spacing. Instead, we found that the N-terminus of Itc1 is essential for proper functioning of the ISW2 complex at ISW2-regulated genes, where ISW2 positions the +1-nucleosome and spaces nucleosomes downstream. With the help of structural predictions, we identified a ‘downWAC’ motif in the N-terminal part of Itc1. This motif forms a structural module together with the WAC motif. The WAC-downWAC module appears to be only loosely associated with the remainder of the complex. It likely helps *Sc*ISW2 to associate with its target genes by functioning as an anchor to DNA or to specific transcription factors.

## Results

We first inspected the domain architecture of *Sc*Itc1. Only four domains have been identified through homology searches in *Sc*Itc1 so far. Among them is an N-terminal WAC domain, a central DDT domain and three WHIM motifs in the C-terminal half (Fig. 1B) [30]. This organization is shared with the orthologues of *Sc*Itc1 in metazoans called ACF1 (aka. BAZ1A in humans), although ACF1 possesses domains C-terminal of the WHIM motifs, including two PHD fingers and a Bromodomain [26].

To discover additional folded regions in *Sc*Itc1, we employed IUPred2 [31]. We identified a region with low disorder (i.e., low IUPred2 score) just C-terminal of the WAC domain. This region is congruent to a motif previously named downWAC [32] (Figs. 1B, C). The protein sequence of the downWAC region appears to be weakly conserved with similar regions in *Hs*ACF1, *Dm*ACF1, and *Hs*BAZ1B, aka. WSTF (Fig. S1A).

As structural data for *Sc*Itc1or of the full *Sc*ISW2 complex are missing, we predicted the structure of the entire ISW2 complex with AlphaFold 3 [33] (Figs. 1A, B, D). The model shows a mostly well-structured, largely α-helical body, with only few disordered regions. Intriguingly, the downWAC region interacts with the WAC domain in the AlphaFold model (Figs. 1D, E). The structural module formed from WAC and downWAC is well conserved across homologous proteins (Figs. 1E, F), suggesting that its function remains conserved as well.

Notably, one side of the WAC-downWAC module possesses a highly positively charged interface (Fig. 1G). Residues from both the WAC and down-WAC regions contribute to this interface (Fig. S1B). The positive charge might allow the module to interact with negatively charged nucleic acids, in line with a possible role in DNA binding [26,27].

We next inspected how the WAC-downWAC module attaches to the remainder of the ISW2 complex. A long linker (aa 272 to 382) connects it to the rest of *Sc*Itc1. This linker is unfolded, as evidenced by a low or very low pLDDT quality score (70 > pLDDT > 50 or pLDDT < 50, respectively), consistent with the IUPred2 prediction above (Fig. 1C).

AlphaFold positions the WAC-downWAC module near the second RecA-like lobe (Lobe2) of Isw2 (Fig. 1D). However, the interaction interface between the module and the rest of Isw2 is small (<600 Å^2^) suggesting an instable interaction or unreliable prediction.

Unstable binding between the WAC-downWAC module and the remainder of the complex is supported by the predicted aligned error (PAE), a measure of how confident the algorithm is in pair-wise distances between amino acids [33,34]. The WAC-downWAC module strongly segregates from the remainder of the complex in the PAE matrix (Fig. 2A). No pairwise distances to any other residue outside of the WAC-downWAC region in *Sc*Itc1 or in *Sc*Isw2 possess high confidence. The same is true for the long, unfolded linker (aa 272 to 382) discussed above. The PAE matrix for an AlphaFold prediction of the orthologous *Dm*ACF complex reveals the same features (Fig. 2B). We therefore propose that the WAC-downWAC modules do not form stable interaction interfaces with the remainder of the complexes in ACF and its orthologues. It is as if the WAC-downWAC modules are attached to the rest of the complexes via a flexible polypeptide linker.

**Fig. 2.**
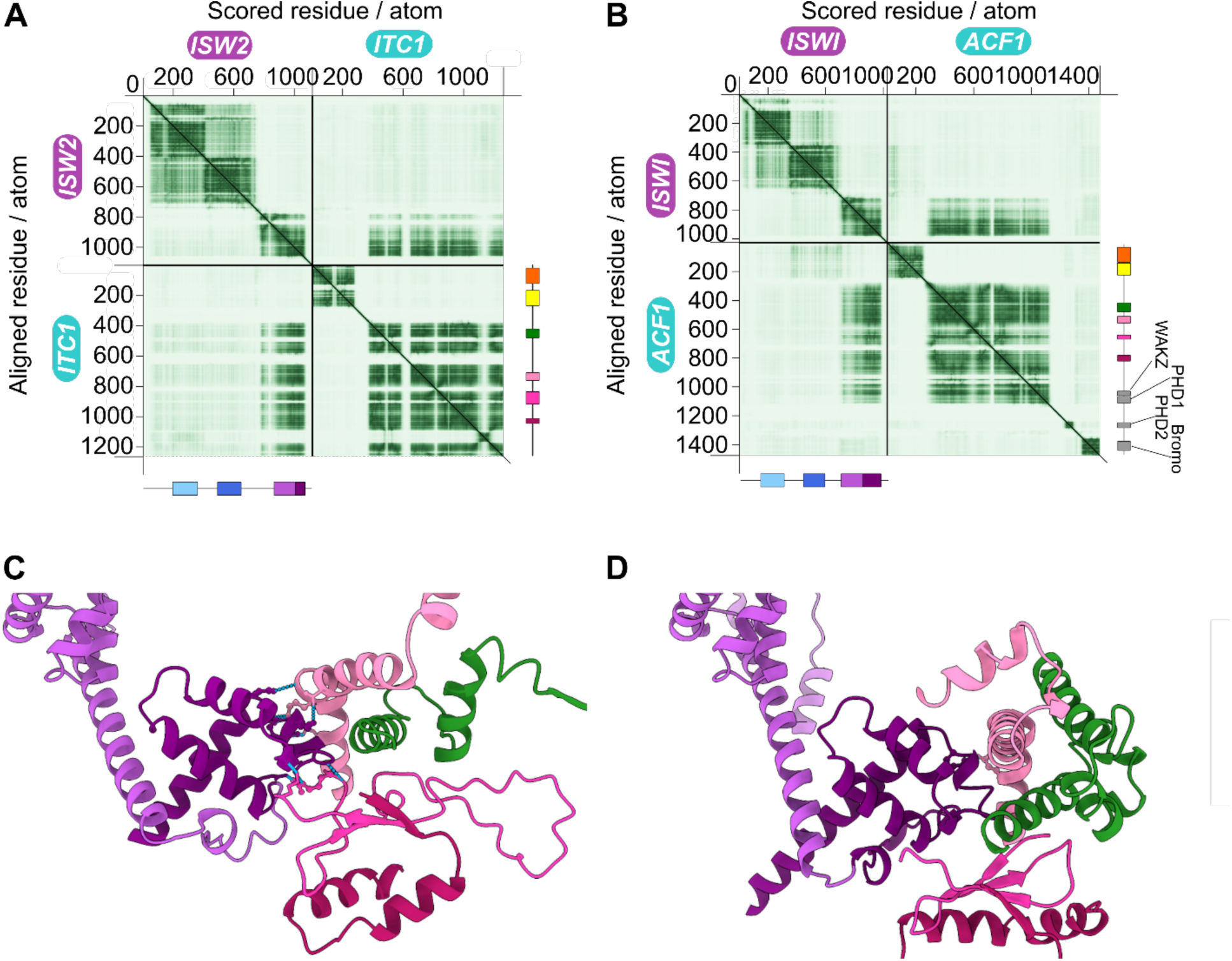
The WAC-downWAC module does not stably associate with other protein domains in the *Sc*ISW2 and *Dm*ACF complexes. **(A)** PAE matrix of the *Sc*ISW2 complex. The N-terminal WAC-downWAC module segregates from the remainder of the matrix. Rigid domain-domain interfaces are predicted to occur between the C-terminal HAND-SANT-SLIDE domain of Isw2 and the body of *Sc*Itc1, including its DDT, WHIM1, WHIM2 and WHIM3 domains. Note the miniature domain organizations on the right and bottom with colors as in Figs. 1A, B. **(B)** PAE matrix of the *Dm*ACF complex. **(C)** The SLIDE domain of *Sc*Isw2 (deep purple) interacts with the WHIM1 (light pink), WHIM2 (pink), WHIM3 (dark pink), and DDT domains (green) of *Sc*Itc1. Interacting amino acids are indicated. **(D)** As panel C, but for *Dm*ACF.

The PAE matrices also hold clues as to how the complexes are held together. The C-terminal quarter of *Sc*Isw2, particularly its SLIDE, has high confidence distance predictions to residues across most regions of *Sc*Itc1 except the N-terminal 382 amino acids. The PAE matrix of the *Dm*ACF structural model conveys the same behavior (Fig. 2B). These results suggest that the subunits rigidly associate and that the SLIDE domain plays a dominant role in this process.

As predicted from the PAE matrices, the SLIDE domains of *Sc*Isw2 and *Dm*ISWI are engulfed by *Sc*Itc1 and *Dm*ACF1. The DDT and the WHIM motifs 1 and 2 contact the tip of the SLIDE domain (Figs. 2C, D) as predicted previously [30].

To investigate the molecular underpinnings of the drastic growth defect caused by deletion of the WAC-downWAC module of *Sc*Itc1 in the BY4741 background [27], we deleted amino acids 2–374 of *Sc*Itc1 in haploid BY4741 cells by direct transformation. We collected three independent colonies and confirmed successful deletion by PCR and whole-locus Sanger sequencing. Surprisingly, we did not observe the previously described growth defects in all three biological replicates (Fig. 3A).

**Fig. 3.**
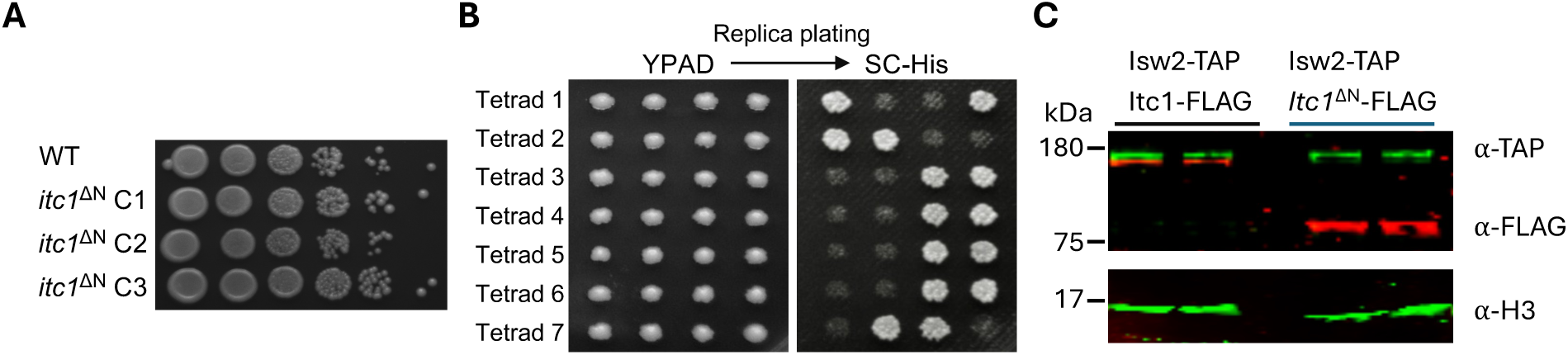
Deletion of the N-terminus of Itc1 does not affect cell viability in *S. cerevisiae* BY4741: **(A)** Spot dilution assay of cells generated by direct replacement of the *ITC1* gene with *itc1*^ΔN^-*HIS3* cassette in the BY4741 background. Ten-fold dilutions are shown. **(B)** Representative tetrad dissection plate of a diploid heterozygous *itc1*^ΔN^/*ITC1* yeast strain. Tetrads were dissected on a YPAD plate, colonies grown for three days and replica-plated on histidine-lacking medium (SC -His) to identify the colonies with the *itc1*^ΔN^-*HIS3* cassette. **(C)** Western blot showing expression of Itc1 and its binding partner Isw2 in whole cell extracts. Proteins were C-terminally tagged with FLAG (Itc1) and TAP (Isw2). Histone H3 served as a loading control. Uncropped images can be found in Fig. S4.

We considered the possibility that our haploid *itc1*^ΔN^ strains only grew normally because they had acquired secondary suppressor mutations that allowed *itc1*^ΔN^ cells to proliferate. We therefore deleted Itc1’s N-terminus in diploid BY4743 cells. We selected heterozygous mutants which harbor one wild-type *ITC1* copy. We then performed sporulation followed by tetrad dissection to form four haploid spores. Two of the four spores should carry the *itc1*^ΔN^ allele. Cells arising from *itc1*^ΔN^ spores would be expected to exhibit severe growth defects because possible suppressor mutations should not strictly segregate with *itc1*^ΔN^. Contrary to the expectation, we observed no growth defects in colonies formed by any of the four spores in full medium (YPDA; 50 tetrads were analyzed from two independently created heterozygous diploids; Fig. 3B, Fig. S2). We confirmed the presence of the *itc1*^ΔN^ allele by replica-plating colonies on minimal medium lacking histidine (SC -His). Moreover, we selected two histidine-prototrophic colonies from the tetrad 1 and confirmed the presence of the *itc1*^ΔN^ allele by PCR and by Sanger sequencing from genomic DNA.

We next considered the possibility that the Itc1^ΔN^ protein is unstable and rapidly degraded, thereby bypassing a toxic gain-of-function phenotype. To test this possibility, we FLAG-tagged the C-terminus of Itc1 in two independent transformants to perform western blots. The western blot results show that cells robustly express *Itc1^ΔN^*-FLAG, even to higher levels than the full-length Itc1-FLAG (Fig. 3C). At the same time, the FLAG-tagged *Itc1^ΔN^* strains showed similar growth as WT cells (Fig. S3). Collectively, the expression analysis strongly argues against the possibility that Itc1^ΔN^ protein is rapidly degraded, thereby masking a toxic phenotype. It remains formally possible though that the C-terminal FLAG tag itself acts as yet another suppressor of toxicity of the N-terminal deletion of Itc1.

Collectively, genetic, phenotypic and expression analysis strongly argue against the possibility that the deletion of the N-terminus of *Sc*Itc1 induces a toxic gain of function phenotype in BY4741 cells. We note that our tetrad dissection may have missed a tightly linked suppressor mutation. This mutation, however, would have had to occur in two independently created heterozygous diploids, which we consider unlikely.

Given the absence of a strong phenotype in *itc1*^ΔN^ cells (this work) and the known specialization of the ISW2 complex to work only on a few hundred genes, we also expected at most mild alterations in the nucleosome landscape. To test this prediction, we performed micrococcal nuclease sequencing (MNase-Seq). As predicted, the genome-averaged nucleosome architecture remained indistinguishable in *itc1*^ΔN^ cells compared to cells with wildtype cells (WT) (Fig. 4A, left, Fig. S5). Also, the average nucleosome-to-nucleosome distance, the so-called nucleosome repeat length (NRL) remained indistinguishable (Fig. 4B left). As controls, we performed full deletions of *ITC1* and *ISW2* (Fig. 4A, left), which also did not lead to noticeable changes compared to WT, confirming earlier work [10,35–37]. We conclude that the *itc1*^ΔN^ mutation does not induce genome-wide changes in nucleosome organization. This result is consistent with the absence of growth phenotype (see above) but cannot be readily reconciled with the previously reported gain of toxic function of this deletion.

**Fig. 4.**
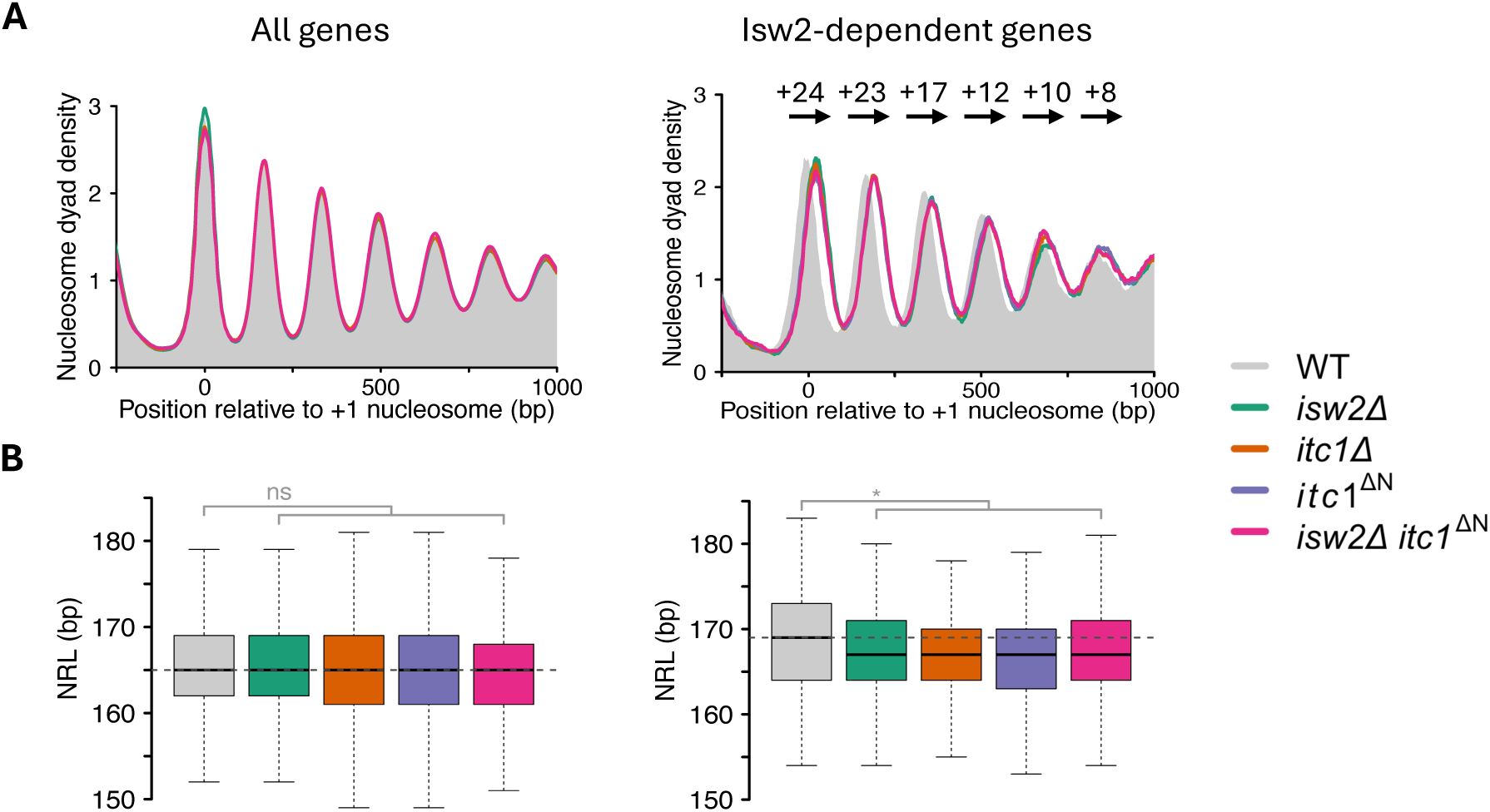
Deletion of the N-terminus of Itc1 protein affects nucleosome organization only at ISW2-specific genes. **(A)** Average nucleosome organization in the indicated yeast strains (see legend). Left: composite nucleosome organization over all yeast genes (5015); right: over Isw2-dependent genes (230; Table S2). A gene was considered to be Isw2-dependent if its +1-nucleosome location was shifted downstream into the gene body by at least 10 bp in *isw2Δ* datasets compared to WT. Values above each peak in left panel indicate a downstream shift of nucleosomes (in bp) in *isw2Δ* mutant relative to WT. **(B)** Boxplots showing nucleosome repeat length (NRL) distribution in the same set of genes as in A. Boxplots represent the median, interquartile range and 1.5X of the interquartile range. Statistical significance comes from a two-tailed Welch’s t-test performed on the mean NRL values of two or three datasets. MNase-Seq was performed twice in *MATa* and once in *MATα* background in all strains except in *itc1*^ΔN^ for which only the *MATa* background was used. Horizontal dotted line indicates the median NRL in WT. See Fig. S5 for the individual mutants overlaid with WT.

The wild-type ISW2 complex is known to slide the +1-nucleosome towards the promoter region at specific genes [18,36]. Therefore, we tested nucleosome organization and NRL at these ISW2-dependent genes. Using DANPOS [38], we identified 230 genes that experienced a downstream shift of their +1-nucleosome by more than 10 bp upon deletion of *ISW2* (Table S1, S2) (Fig. 5). The +1 nucleosomes of these genes shifted by the same amount, 24 bp, in *itc1Δ* and *itc1*^ΔN^ cells (Fig. 4A right; Table S1), suggesting that the mutant ISW2 complexes are similarly defective in +1-nucleosome positioning as cells lacking the Isw2 ATPase subunit.

**Fig. 5.**
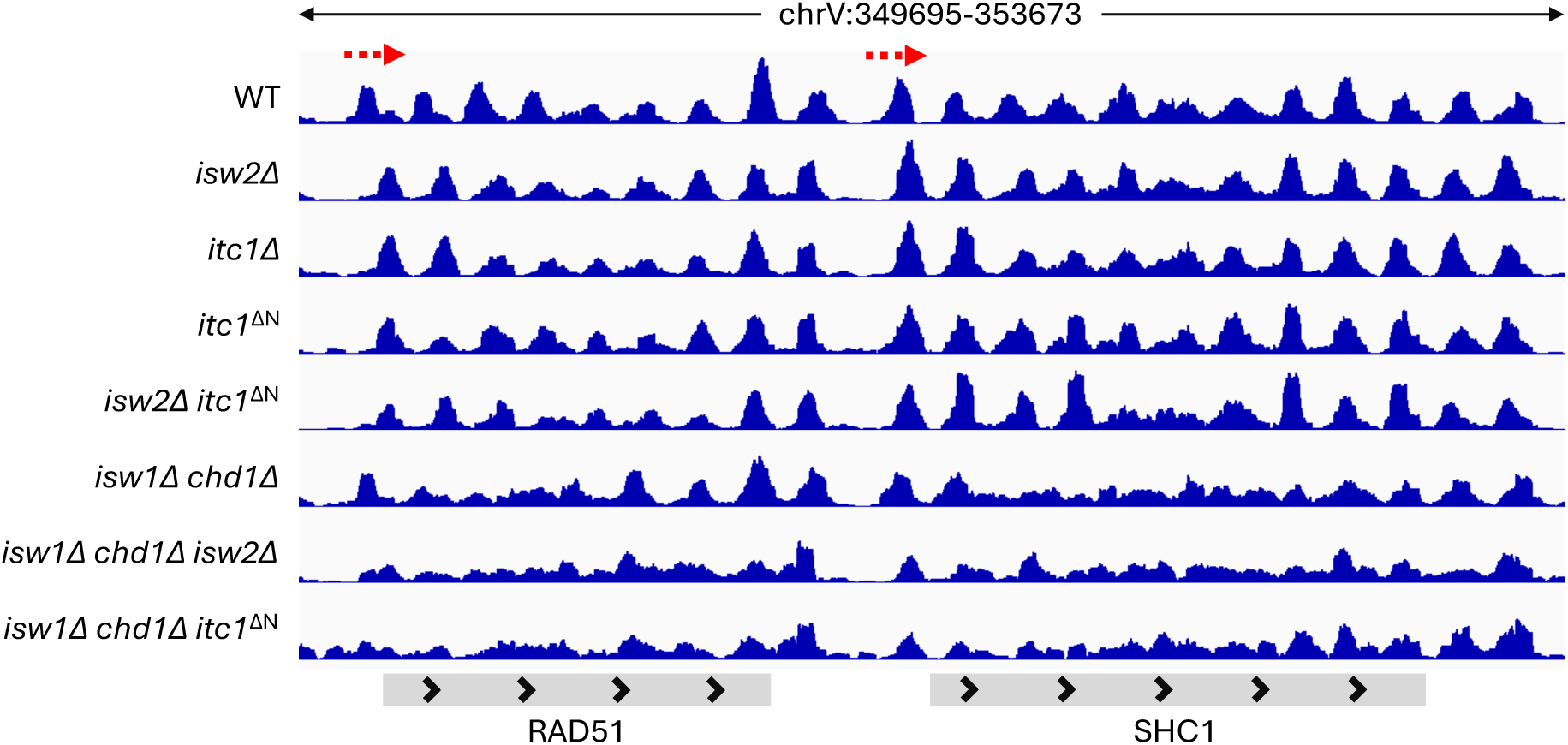
Genome browser snapshot over two representative genes in the indicated yeast strains. Red dashed arrows represent the +1-nucleosomes that experience a downstream shift in *itc1* and *isw2* mutant cells.

Nevertheless, we found that the NRL in ISW2-dependent genes (Table S2) in *isw2Δ* cells is decreased by 2 bp compared to wild-type (WT), possibly arising from the 3’ shifted promoter-proximal nucleosomes. The NRL over the same genes is also decreased by 2 bp in *itc1Δ* cells and even in *itc1*^ΔN^ cells (Fig. 4B right). The results suggest that deletion of Itc1’s N-terminus does not allow the ISW2 complex to function properly *in vivo*. Furthermore, the double deletion of Isw2 ATPase and the Itc1’s N-terminus did not lead to additive effects, confirming that the phenotype seen in *itc1Δ* cells is mediated through Isw2 (Figs. 4A, 4B, 5). Lastly, we tested the effects of deleting just the WAC motif in available MNase-Seq datasets [28] in the W303 background (Fig. S6). The *itc1*^ΔWAC^ phenocopied the full deletion of *isw2*, supporting the idea that WAC and downWAC form a structural and functional module.

We next considered the possibility that effects of N-terminal ablation of Itc1 on the growth phenotype and on the nucleosome architecture may be obscured by other remodeling factors. In fact, the ISW1 and Chd1 remodelers can act redundantly to ISW2 on the nucleosome organization at shared genes [36,37]. Therefore, we studied the effects of the *itc1*^ΔN^ allele in *isw1Δ chd1Δ* double knockout (DKO) cells. No growth defect became apparent upon deletion of Itc1’s N-terminus, nor did the loss of the *ISW2* gene cause a growth phenotype (Fig. 6A). Also the overall chromatin architecture was unchanged compared to DKO cells (Fig. 6B left; Figs. S7, S8). The results again do not support a gain of toxic function phenotype of the *itc1*^ΔN^ allele even in absence of two remodelers that could potentially obscure molecular phenotypes through functional redundancy.

**Fig. 6.**
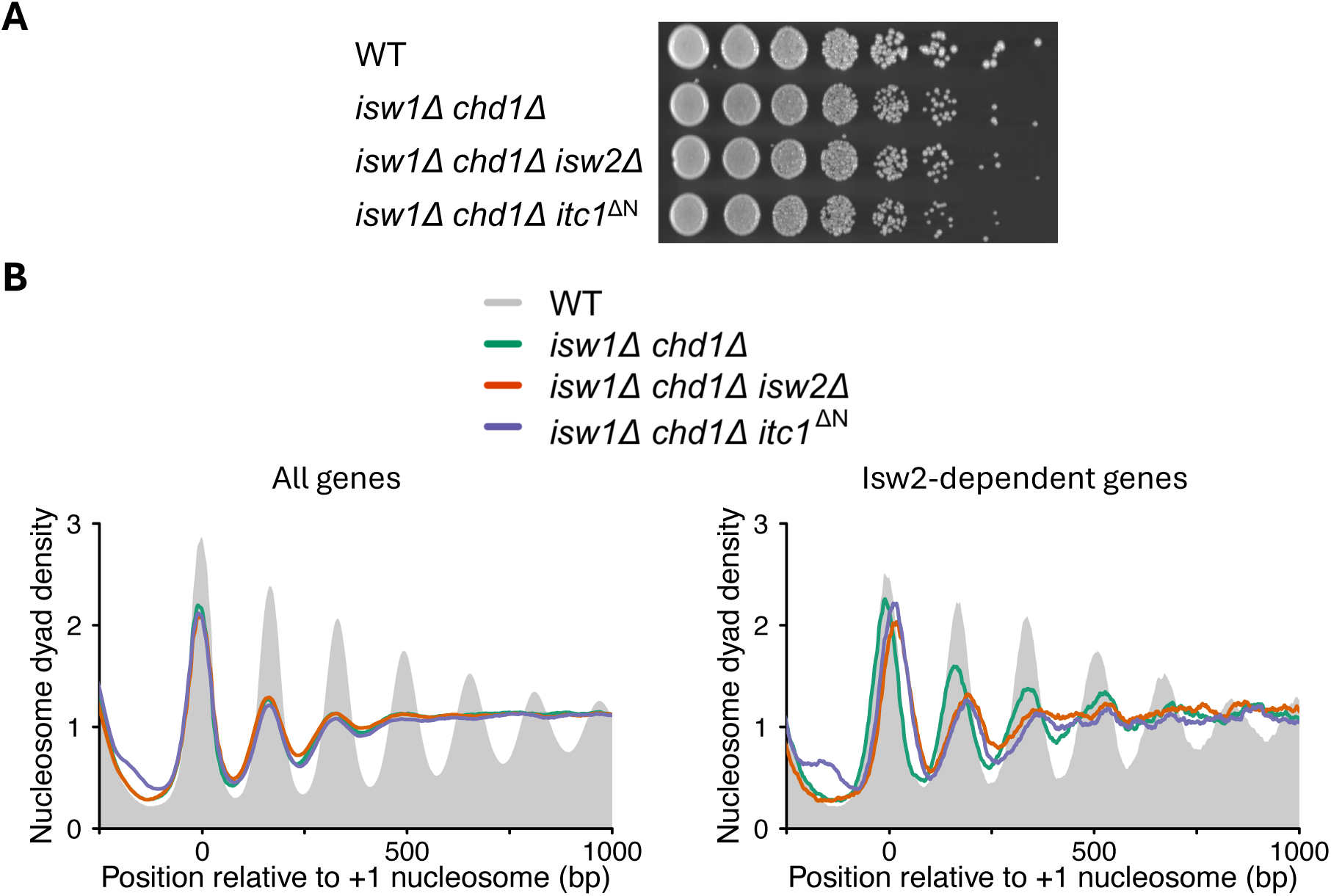
Deletion of the N-terminus of Itc1 does not affect cell viability and nucleosome organization in *isw1Δ chd1Δ* double knockout cells. **(A)** Spot dilution (five-fold) assay of cells generated by direct replacement of Itc1 gene with the N-terminus lacking Itc1 construct in *isw1Δ chd1Δ* background. **(B)** Composite plots showing nucleosome organization of all yeast genes (left) and genes exhibiting a shift in the +1- nucleosome location by at least 10 bp in *isw2Δ* strain (right) in the indicated yeast strains. MNase-Seq was performed once in *MATa* and *MATα* background for mutant strains with similar results. Shown are data from the *MATa* background. See Fig. S7 for deletions in *MATα* background and Fig. S8 for individual mutants overlaid with WT sample.

Finally, we tested if ISW2 needs the activity of Isw1 and Chd1 remodelers to function properly, and if Itc1’s N-terminus is needed for this activity. The +1 nucleosomes on ISW2-dependent genes are similarly downstream-shifted in *itc1*^ΔN^ and *isw2*Δ cells compared to DKO cells (Figs. 5, 6B right; Figs. S7, S8). The magnitude of the shift (23 bp and 25 bp in *itc1*^ΔN^ and *isw2*Δ-deleted DKO cells, respectively) is comparable to the downstream shift seen in cells with functional Isw1 and Chd1 remodelers (Fig. 4A right, Table S1). The lack of well-positioned nucleosomes in the gene bodies in DKO cells prevented us from calculating NRL at individual genes (Fig. 5). We conclude that the ISW2 remodeling complex does not depend on Chd1 and Isw1 remodeling factors to position the +1 nucleosomes, and that the ablation of ITC1’s N-terminus phenocopies the deletion of *ISW2*.

## Discussion

ISWI is a highly conserved ATPase that forms a diverse set of remodeling complexes in eukaryotes [39,40]. The functions of associating subunits, like that of Acf1 in ACF/CHRAC or the orthologous Itc1 in the yeast ISW2 complex, are poorly understood. Previous work implicated the N-terminal region of human Acf1 in linker length sensing, implying that it is functionally needed to space nucleosomes [27]. The reported drastic growth phenotype associated with the *itc1*^ΔN^ allele underscored the importance of the N-terminus. It also implied that the mutant ISW2 complex has gained a toxic function, perhaps by producing irregular spacing if nucleosomes in parts of the genome.

Unexpectedly, we found Itc1’s N-terminus including the WAC domain to be completely dispensable for growth in BY4741 cells. We searched for but did not find possible suppressor mutations in our mutant strains that would reconcile our data with the previously reported drastic growth defect. It remains possible, however, that the introduction of our selection cassette (*HIS3*) inadvertently introduced a suppressor. CRISPR-guided gene editing could be used in the future to surgically remove Itc1’s N-terminus without leaving other traces in the genome.

Loss of the Itc1 N-terminus also did not induce rogue remodeling of nucleosomes as might be expected if from a toxic remodeler mutation. Collectively, our results do not support a gain of toxic function phenotype. Given that Itc1 N-terminus is unessential in W303 yeast cells as well [28], we propose that loss of the Itc1’s N-terminal region is generally well tolerated under standard growth conditions in *S. cerevisiae*.

We find instead that deletion of Itc1’s N-terminus leads to a full ISW2 loss of function phenotype. In none of our assays could we distinguish *itc1*^ΔN^ from *isw2*Δ or *itc1*Δ alleles. These results are in line with a recent study that proposes that ISW2 relies on Itc1’s N-terminus to identify its target genes [28]. In fact, we found similar set of genes showing the +1-nucleosome shift upon deletion of the Isw2 ATPase, Itc1 or the N-terminus of Itc1 (Table S2). This result argues against the possibility that the mutant ISW2 complex is mistargeted to a different set of genes than WT and instead points to a loss of function phenotype associated with the *itc1*^ΔN^ allele.

Our structural predictions suggest that the WAC-downWAC region forms a structural module that is flexibly linked to the remainder of the complex. The WAC domain contains a transcription factor binding motif [28], and it associates with DNA [27,30]. We therefore speculate that the WAC-downWAC module may help the remodeling complex to scan the genome for cognate transcription factors (TF). Once the module latches onto a TF, the long and flexible polypeptide linker may act like a ‘leash’ for the remainder of the complex, allowing ISW2 to explore the immediate vicinity for nucleosomes that need to be translocated (Fig. 7).

**Fig. 7.**
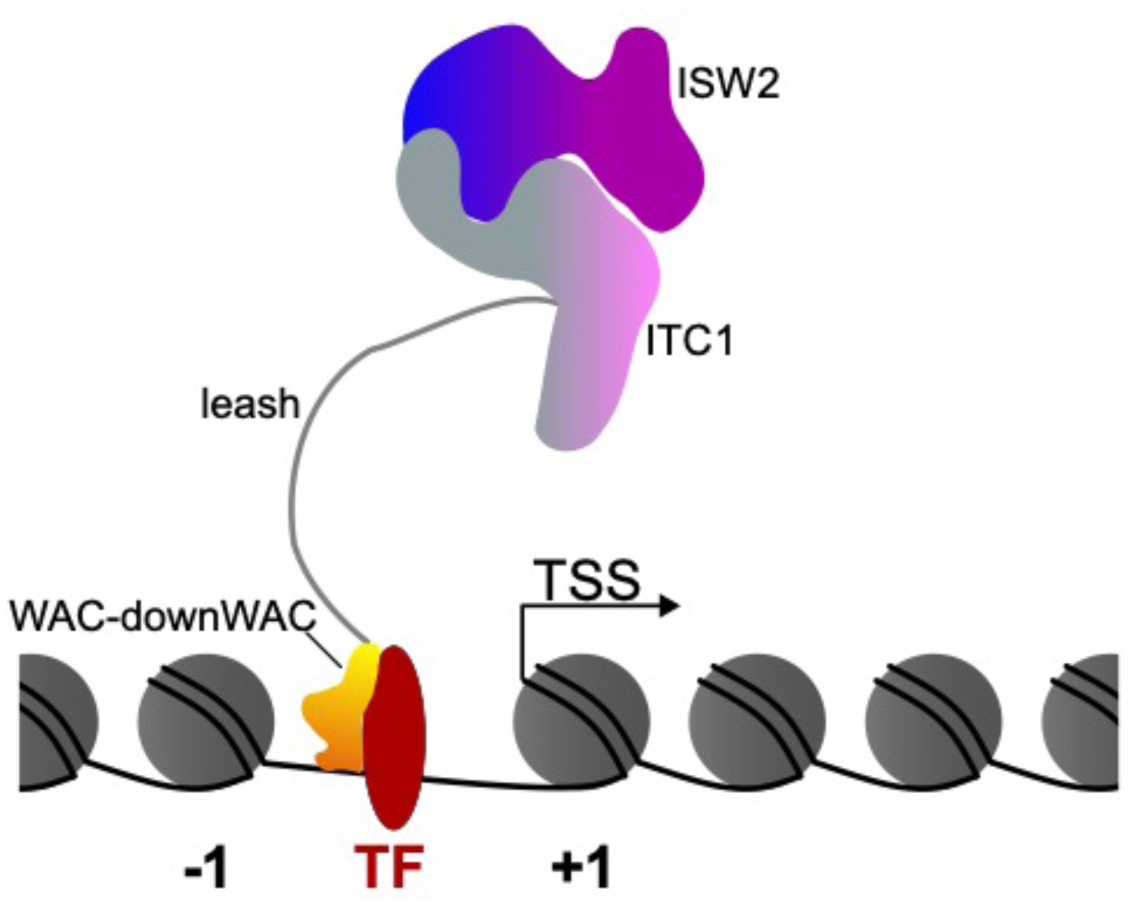
Speculative Model. The WAC-downWAC module (yellow-orange) is connected to the remainder of the complex (blue, purple, grey; colors similar to Figs. 1A, B, D) via a long unstructured polypeptide linker. This linker may facilitate the WAC-downWAC module to scan the genome for appropriate transcription factors (TF, red) [28]. It then anchors the complex near the TF-bound DNA [26,27]. The long linker allows remodeling of nucleosomes that are in the immediate vicinity.

## Methods

### Structural predictions

AlphaFold 3 [33] was used to predict the structures of *Sc*Itc1 (Uniprot ID: P53125) bound to *Sc*Isw2 (Uniprot ID: Q08773), of *Hs*ACF1 (Uniprot ID: Q9NRL2), of *Dm*ACF1 (Uniprot ID: Q9V9T4), and of *Hs*BAZ1B (Uniprot ID: Q9UIG0). All structures were aligned to the WAC-downWAC domain of *Sc*Itc1 using the matchmaker tool in ChimeraX [41]. The RMSD values were calculated relative to *Hs*ACF1. PAE matrices were generated using PAE viewer [34]. The protein disorder prediction of *Sc*Itc1 was generated using the long option of IUPred2A [31]. Figures were designed using Inkscape Version 1.2.

### Yeast strain generation

All yeast strains used in the study are derived from the S288c background (Table S3). To delete one copy of the N-terminus of Itc1, the *ITC1* gene was first cloned in pRS406 plasmid with the *HIS3* marker gene downstream of the Itc1 gene to yield plasmid pFMP773. Amino acids 2-374 of Itc1 were deleted from pFMP773 using oligonucleotides oFMP722 and oFMP723 by Gibson cloning (Table S4), yielding pFMP777. The insert was fully sequenced with Sanger sequencing (oFMP513 – oFMP518 and oFMP572 – oFMP575). The mutant construct was PCR amplified with oFMP544 and oFMP773, agarose gel purified and transformed into BY4743 (yFMP721) strain. Cells were plated on SC-His media and single colonies were picked after three days. Genomic DNA was isolated and the *itc1* locus was tested by PCR using oFMP456, oFMP457 and oFMP502. Colonies that showed two bands corresponding to the WT *ITC1* and the mutant allele were selected, yielding yFMP298 and yFMP299.

For direct deletion of the N-terminus of Itc1 in haploid yeasts, the mutant construct along with *HIS3* gene was PCR-amplified using oFMP544 and oFMP773 from pFMP773 and transformed into BY4741 (yFMP009) strain, yielding yFMP039, yFMP040 and yFMP041. The mutant strains were fully sequenced with Sanger sequencing for any mutations at the Itc1 locus using oFMP513 – oFMP518 and oFMP572 – oFMP575. The WT or mutant Itc1 genes were C-terminally FLAG-tagged as follows: The plasmids pFMP773 and pFMP777 were PCR amplified using oligos oFMP700 and oFMP701. The FLAG-tag sequence was PCR amplified from the plasmid pFMP741 with oligos oFMP702 and oFMP703. Both PCR fragments were assembled using Gibson assembly to C-terminally tag the WT or mutant Itc1 genes leading to plasmids pFMP775 and pFMP757, respectively. As above, the FLAG-containing constructs were PCR amplified with oFMP544 and oFMP773, agarose gel purified and transformed into yFMP168, yielding yFMP197, yFMP198, yFMP200 and yFMP202. The successful insertion at the Itc1 locus was confirmed by PCR.

To generate yeast strains lacking combinations of genes *ISW1*, *CHD1*, *ISW2* and the N-terminus of Itc1, first cells lacking *ISW1* and *CHD1* in opposite mating type (yFMP001 and yFMP296) were mated (yFMP294). Yeast strain lacking both *ISW1* and *CHD1* were generated by sporulation and tetrad dissection (yFMP360, 362). To generate a triple mutant of *ISW1*, *CHD1* and *ISW2* or N-terminus lacking Itc1, yFMP362 was mated strain lacking *ISW2* (yFMP007) or the N-terminus of Itc1 (yFMP039), followed by sporulation and tetrad dissection. Loci *ISW1* and *ISW2* with the same selection marker (KanMX) were confirmed by PCR (oFMP628, 629, 671, 672).

### Sporulation and tetrad dissection

Diploids lacking one copy of the N-terminus of Itc1 were grown in YPA + 4% glucose media and sporulated in minimum sporulation media lacking Histidine (1% KOAc, leucine 25 mg/L, uracil 5 mg/L) at 23 C° for 4 – 5 days. Spores were dissected on a Singer MSM 400 dissection microscope on YPAD plates and grown at 30 C° for 2-3 days. Tetrads were replica plated on SC-His plate and grown for 1-2 days.

### Growth assay

Cells were grown overnight in YPAD media and diluted to OD 600 1.0 in sterile water. 10- or 5-fold dilutions were generated and 10 μl were plated on YPAD media. Images were taken after 3 days of incubation at 30 °C. The assay was performed twice.

### Western blot

Whole cell extracts of cells grown in YPAD media to OD 0.8 were prepared using the NaOH/TCA precipitation method [11]. Blots were blocked with PBS + 5% skimmed milk + 0.1% Tween-20. The primary antibodies used were anti-FLAG (Sigma Cat# F1804, 1:20,000 dilution) and anti-H3 (Abcam Cat# ab1791, 1:20,000 dilution). Western blot was performed twice.

### MNase-Seq

Yeast nuclei preparation, micrococcal nuclease digestion and high-throughput sequencing libraries preparation were done as previously described [11,42]. Briefly, cells were grown in 500 ml YPAD media to OD 600 1.0. Cells were harvested by centrifugation (3000 x g, 8 min, 4 C°), washed with cold water and resuspended in 0.7 M ß-mercaptoethanol, 28 mM EDTA pH 8.0. Cells were incubated at 30 C° for 30 min, washed with 40 ml ice-cold 1 M sorbitol and resuspended in 5 volumes of 1 M sorbitol, 5 mM ß-mercaptoethanol. 2 mg Zymolyase was added and incubated at 30 °C for 20 min. Spheroplasts were harvested (2500 x g, 5 min, 4 °C) and washed with 40 ml ice-cold 1 M sorbitol. Pellets were dissolved in Ficoll buffer (18% Ficoll, 20 mM KH_2_PO_4_ pH 6.8, 1 mM MgCl_2_, 0.25 mM EGTA, 0.25 mM EDTA) and nuclei were aliquoted by centrifugation (12000 x g, 30 min, 4 °C). Nuclei were washed with MNase digestion buffer (15 mM Tris-Cl pH 7.5, 50 mM NaCl, 1.4 mM CaCl_2_, 0.2 mM EGTA pH 8.0, 0.2 mM EDTA pH 8.0, 5 mM ß-mercaptoethanol) and incubated with different amounts of MNase (4 – 256 U / ml) for 20 min at 37 °C. MNase digestion was stopped by adding 10 mM EDTA, 1% SDS and 50 mM Tris-Cl pH 8.0. DNA was isolated with Proteinase K digestion, followed by Phenol: Chloroform: Isoamyl alcohol extraction. RNA was removed by incubation with 10 μg RNase A at 37 °C for 30 min. DNA was ethanol precipitated and resolved on a low-melt agarose gel to test the digestion degree. Samples with 70 % mononucleosome band were chosen for library preparation using the NEBNext Ultra II DNA Library Prep kit for Illumina. Libraries were sequenced for 50 cycles in the paired-end mode.

### Data analyses

All MNase-Seq data were analyzed as described previously [11]. Briefly, fastq files were mapped to *S. cerevisiae* sacCer3 R64-1-1 genome using Bowtie2 v2.2.9 with default settings, except -X 500, -no-discordant, -no-mixed options [43]. Reads mapping to the rDNA region (chrXII 451000:469000) were removed and nucleosome dyad coverage files were generated with fragment lengths 140 – 160 bp. The nucleosome dyad was extended by 50 bp to generate bigwig files. The nucleosome repeat length of each gene were calculated as described before [11] after subsampling all datasets to 5 million reads (https://github.com/musikutiv/tsTools) that adapts [10]. To identify the +1 nucleosome, the dpos command in DANPOS2 with parameters -jw 5, -q 200, -m 1 was used [38]. Genes showing +1 nucleosome shift in all (two or three) available replicates were considered.

## Declarations

**Ethics approval and consent to participate:** Not Applicable

**Consent for publication:** Not Applicable

**Availability of data and materials:** All high-throughput sequencing data have been deposited at the Gene Expression Omnibus under the accession number GSE205956. **Competing interests:** Not Applicable

## Acknowledgements

We thank Răzvan V. Chereji and David J. Clark for sharing NRL calculation scripts, Stefan Krebs and Helmut Blum (LAFUGA, Gene Center, LMU Munich) for sequencing, Tamas Schauer and Tobias Straub for bioinformatics help. We are also grateful to Sigurd Braun and the Department of Physiological Chemistry, Biomedical Center, LMU Munich for access to the tetrad dissection microscope. A.K.S acknowledges support from the Deutscher Akademischer Austauschdienst for a predoctoral scholarship.

## Author Contributions

Ashish Kumar Singh: Conceptualization, Data curation, Formal analysis, Investigation, Methodology, Validation, Visualization, Writing – original draft, Writing – review and editing.

Sabine Ines Grünert: Formal analysis, Investigation, Validation, Visualization, Writing – review and editing.

Lena Pfaller: Formal analysis, Investigation.

Felix Mueller-Planitz: Conceptualization, Methodology, Investigation, Funding acquisition, Project administration, Supervision, Writing – original draft, Writing – review and editing.

---

**Fig. S1.**
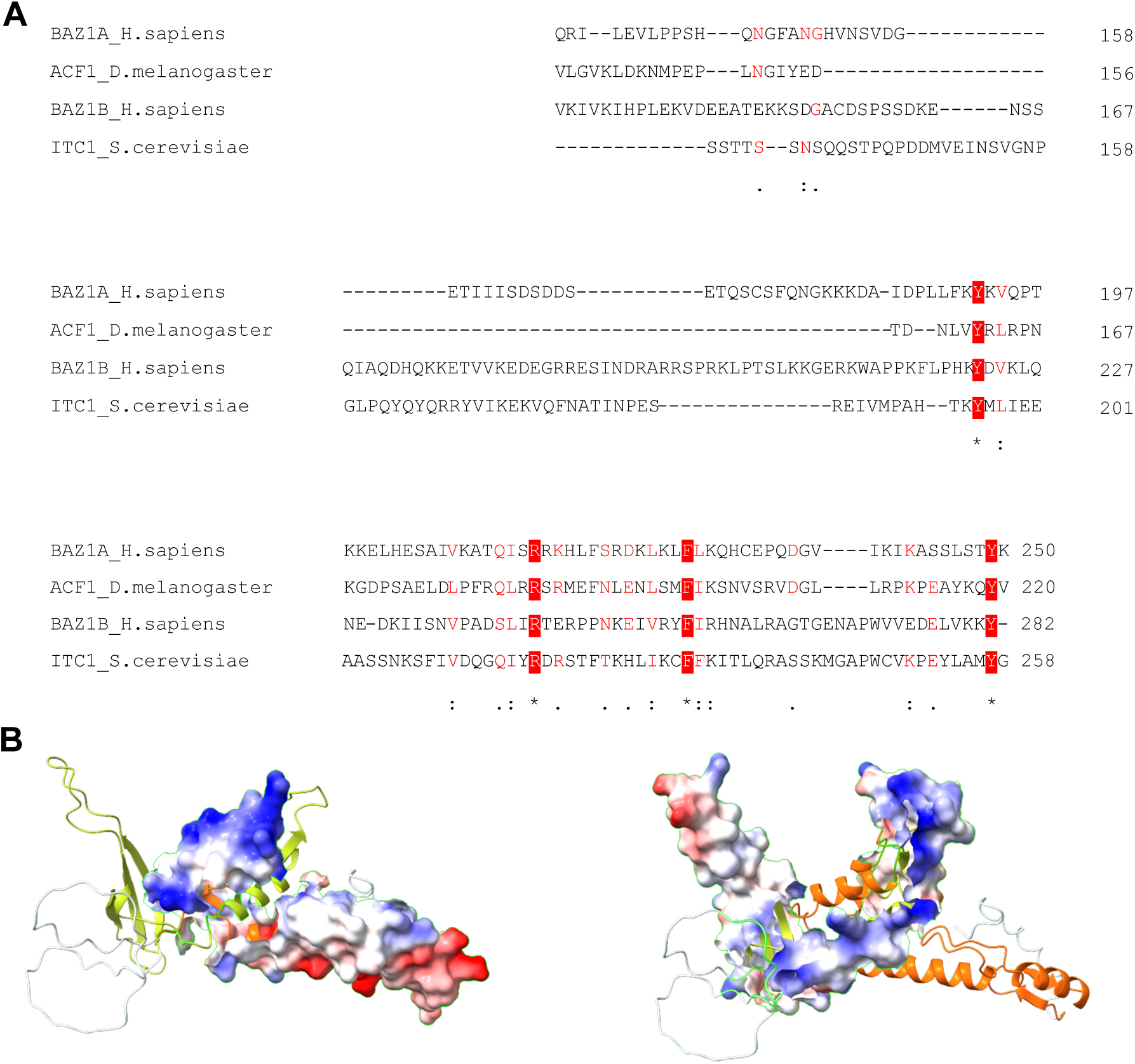
Sequence and structure of the WAC-downWAC module. **(A)** Sequence homology of the downWAC region. **(B)** WAC and downWAC jointly form a positively charged interface. Electrostatic surface potentials of the WAC (left; downWAC is yellow) and downWAC region (right; WAC is orange).

**Fig. S2.**
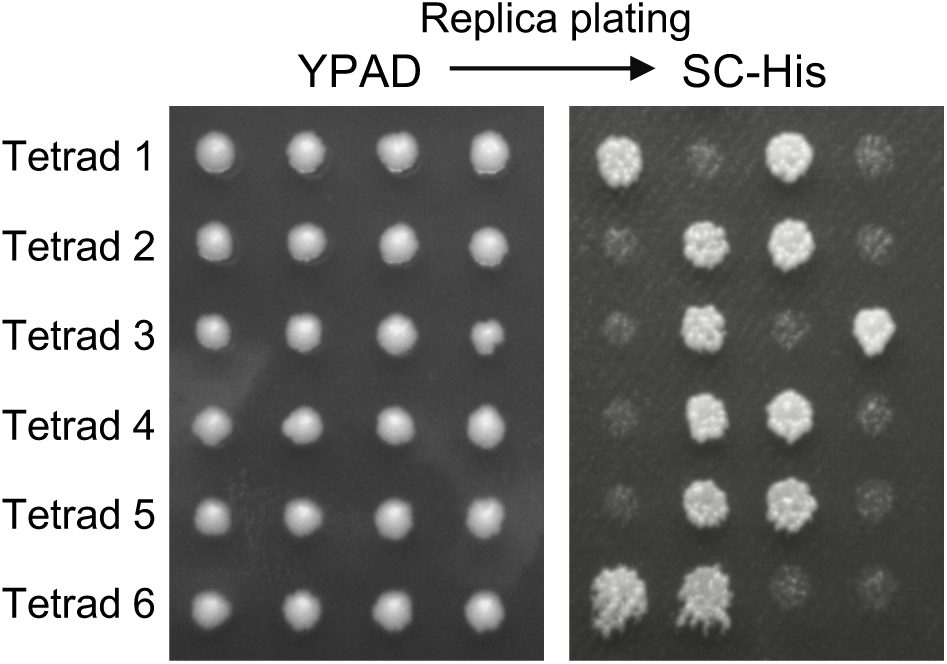
Representative tetrad dissection results from an independent experiment.

**Fig. S3.**
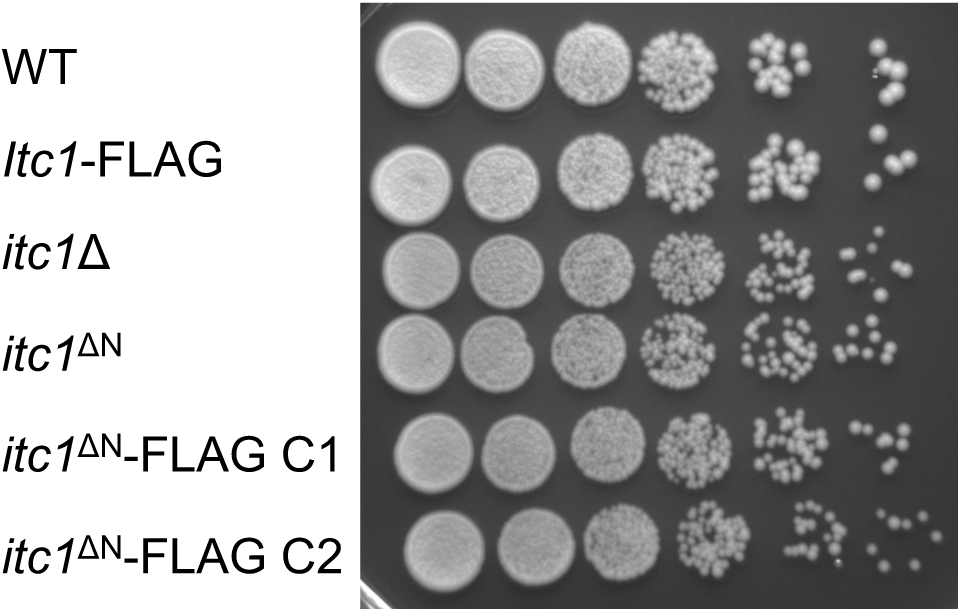
Spot dilution assay of cells lacking Itc1 or with different variants of Itc1 with or without the FLAG tag. Ten-fold dilutions are shown.

**Fig. S4.**
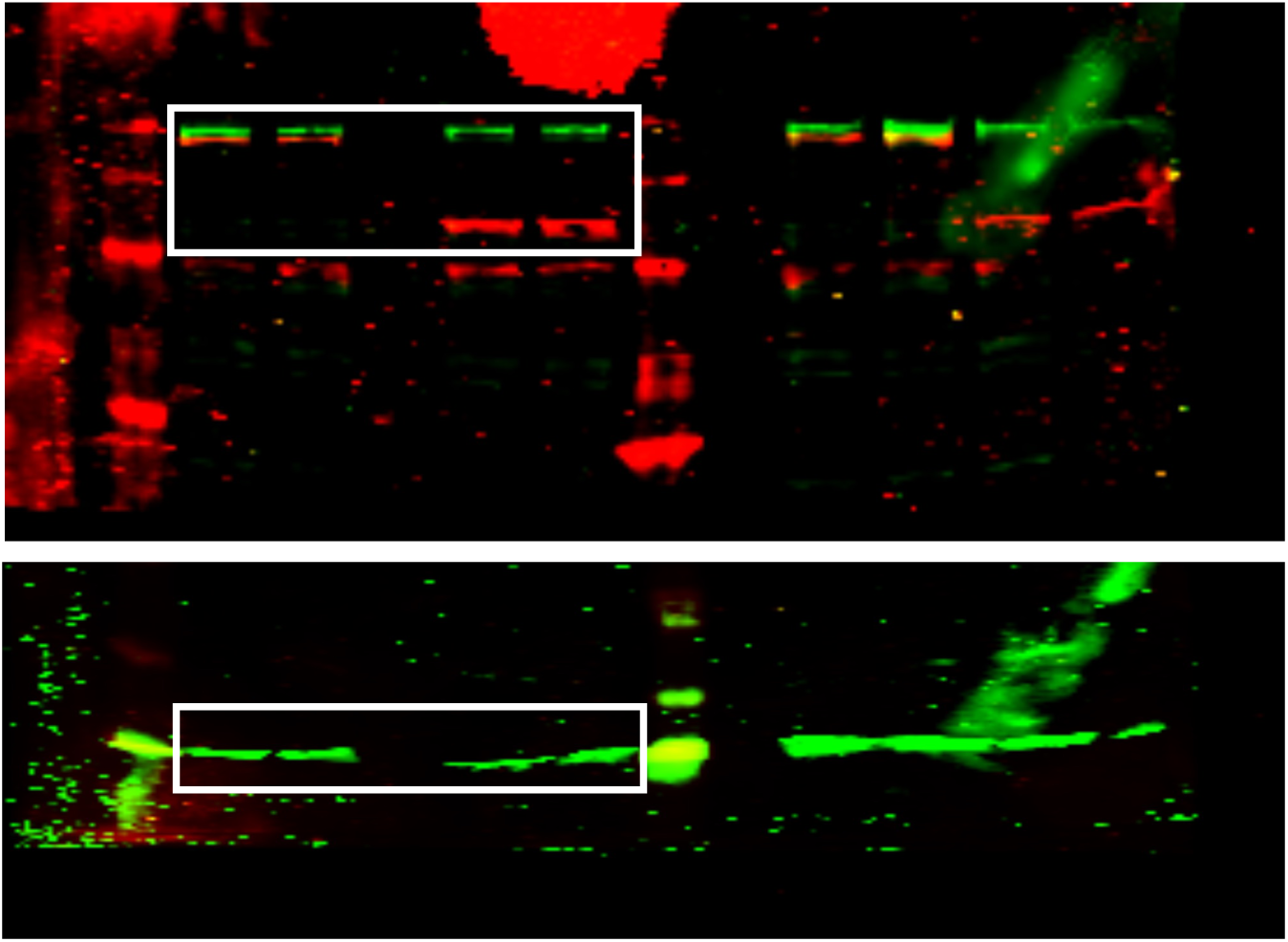
Uncropped blot (related to Fig. 3C).

**Fig. S5.**
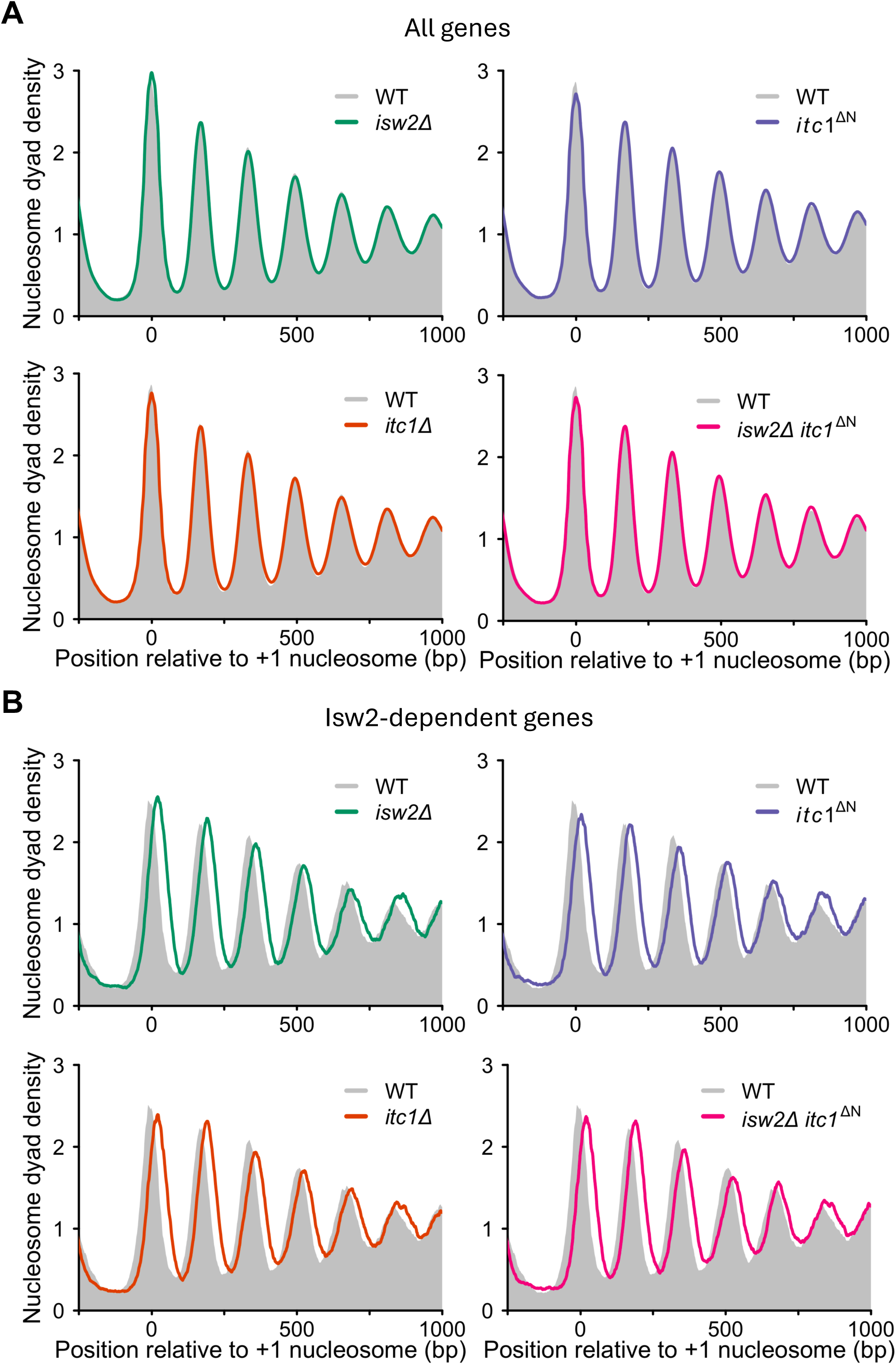
Individual composite plots of indicated yeast mutants overlapped with WT sample for all samples shown in. Fig. 4A**. (A)** Composite plot for all genes. **(B)** Same as A but for Isw2-dependent genes.

**Fig. S6.**
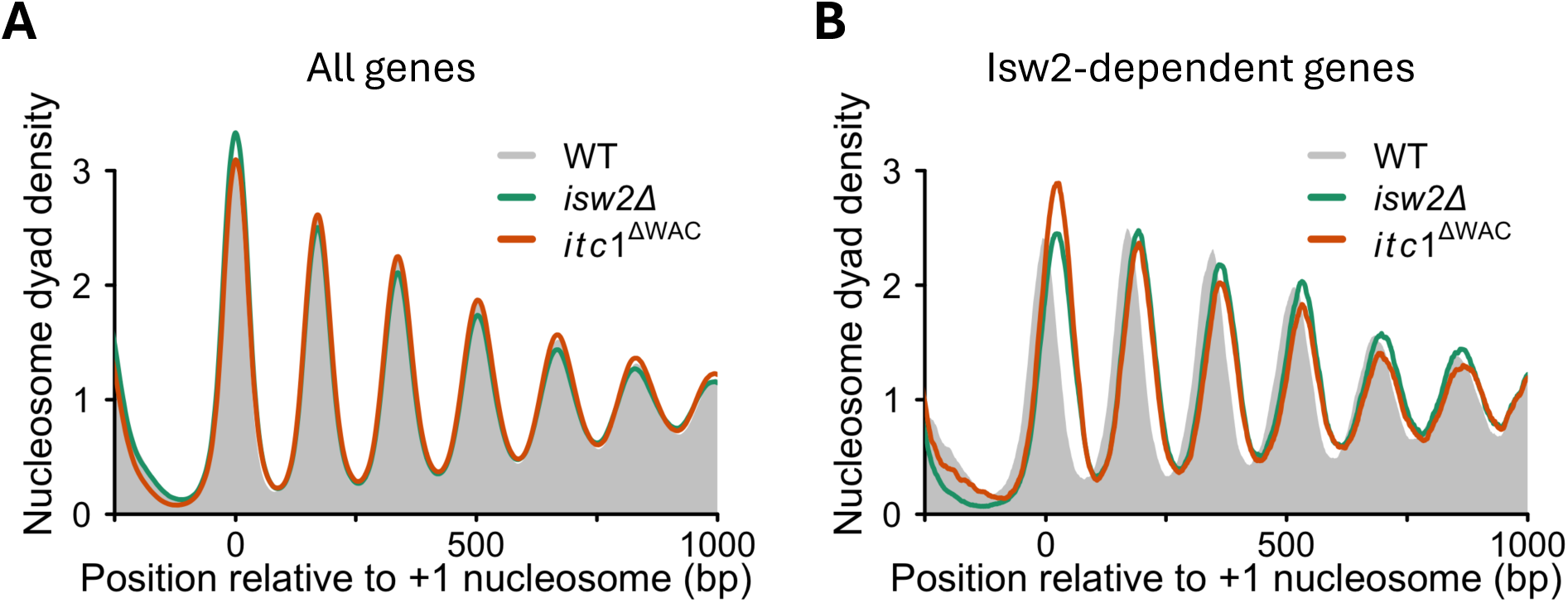
Composite plots of WT and indicated yeast mutants in W303 cells from Donovan et al study. **(A)** Composite plot for all genes. **(B)** Same as A but for Isw2-dependent genes. The *itc1*^ΔWAC^ refers to deletion of amino acids 24-130.

**Fig. S7.**
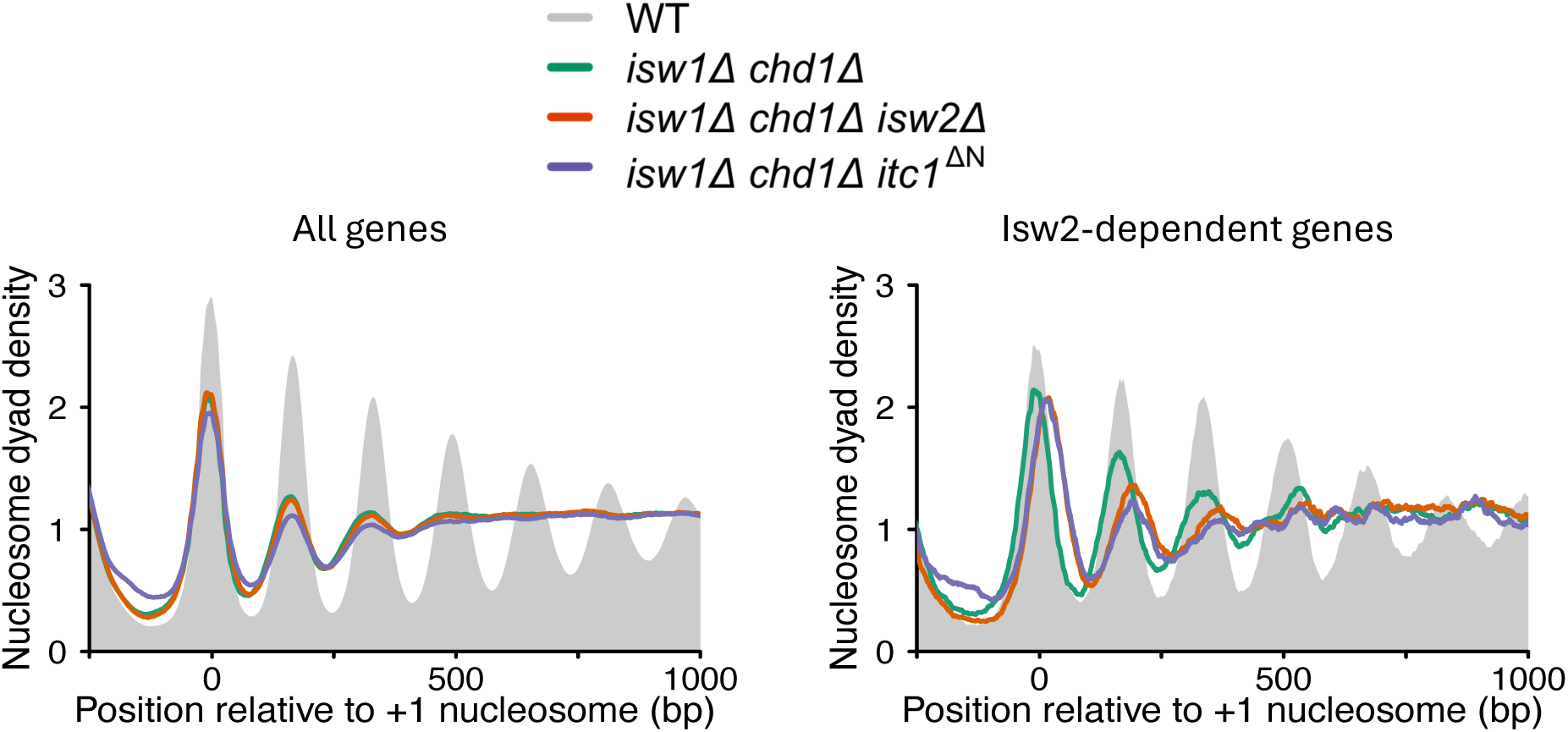
Deletion of the N-terminus of Itc1 does not affect nucleosome organization in *MATα isw1Δ chd1Δ* sensitive background. Composite plots showing nucleosome organization of all yeast genes (left) and genes exhibiting a shift in the +1-nucleosome location by at least 10 bp in *isw2Δ* strain (right) in the indicated yeast strains in *MATα* (alpha) background.

**Fig. S8.**
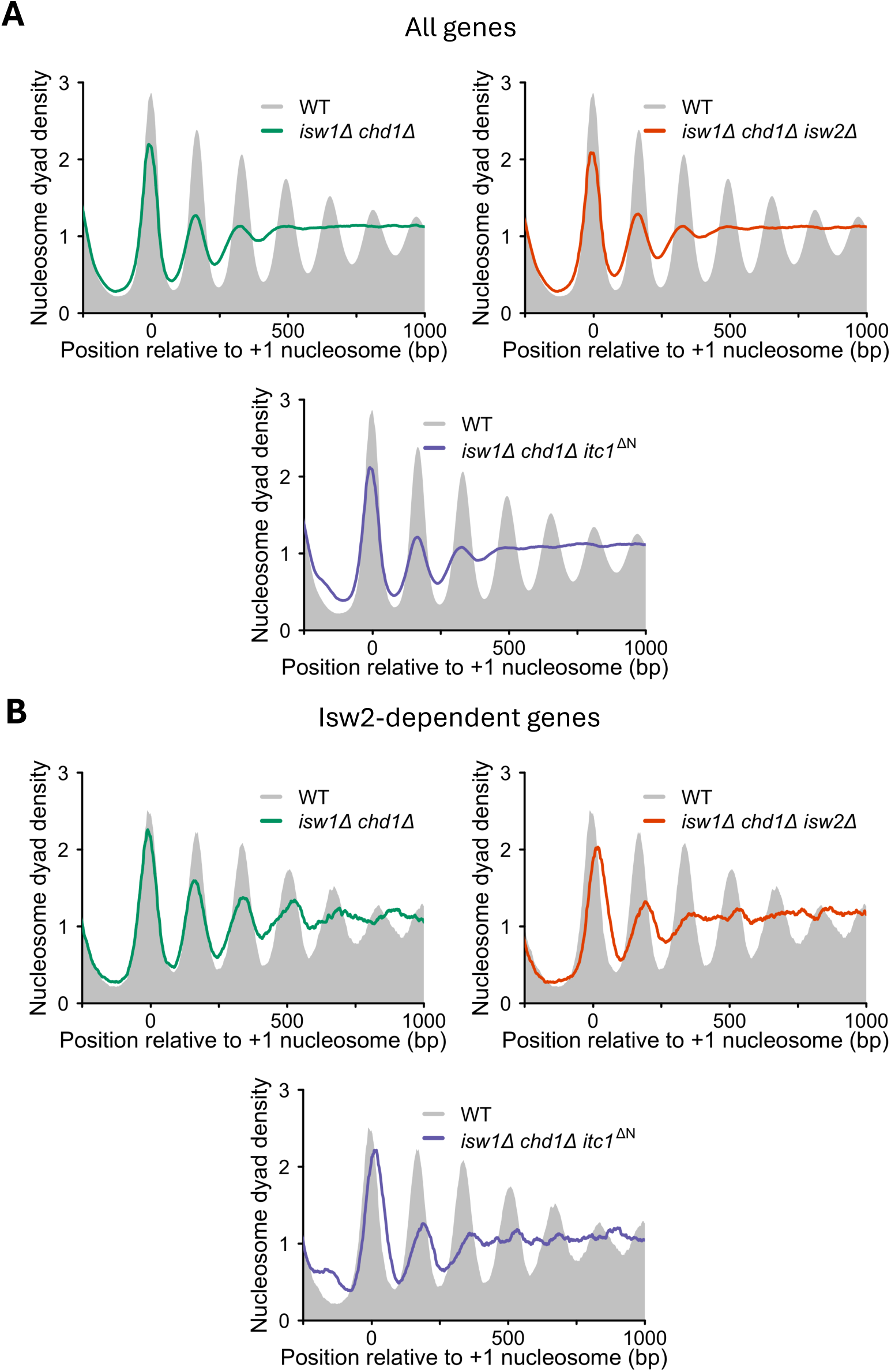
Individual composite plots of indicated yeast mutants overlapped with WT sample for all samples shown in Fig. 6B. **(A)** Composite plot for all genes. **(B)** Same as A but for Isw2-dependent genes.

**Table S1:** Quantification of nucleosome shift in mutant strains relative to WT based on composite plots. Related to Fig. 4A. Positive values indicate a downstream shift and negative values indicate an upstream shift relative to the corresponding WT nucleosome in the composite plot. “NA” indicates cases when a clear nucleosome peak is not identified due to extremely fuzzy nucleosomes in the composite plots.

|  | Nuc +1 | Nuc +2 | Nuc +3 | Nuc +4 | Nuc +5 | Nuc +6 |
| --- | --- | --- | --- | --- | --- | --- |
| *isw2Δ* | +24 | +23 | +17 | +12 | +10 | +8 |
| *itc1Δ* | +24 | +22 | +18 | +13 | +9 | +6 |
| *itc1ΔN* | +24 | +23 | +18 | +11 | +10 | +6 |
| *isw2Δ itc1ΔN* | +25 | +24 | +16 | +12 | +8 | +5 |
| *isw1Δ chd1Δ* | -4 | +2 | +5 | +15 | NA | NA |
| *isw1Δ chd1Δ isw2Δ* | +25 | +23 | +21 | +25 | NA | NA |
| *isw1Δ chd1Δ itc1ΔN* | +23 | +22 | +22 | +25 | NA | NA |

**Table S2:** List of genes showing +1 nucleosome shift by at least 10 bp in indicated yeast strains compared to WT.

|  |  |  |  |  |  |  |
| --- | --- | --- | --- | --- | --- | --- |
| *isw2Δ* | *itc1Δ* | *itc1<sup>ΔN</sup>* | *isw2Δ itc1<sup>ΔN</sup>* | *isw1Δ chd1Δ* | *isw1Δ chd1Δ isw2Δ* | *isw1Δ chd1Δ itc1<sup>ΔN</sup>* |
| YAL018C | YAL018C | YAL018C | YAL002W | YAL042W | YAL042W | YAL042W |
| YAL043C | YAL043C | YAL043C | YAL018C | YBL004W | YAL043C | YAL043C |
| YAR015W | YAL060W | YAR018C | YAL043C | YBR085W | YAL054C | YAR015W |
| YAR018C | YAR018C | YBL023C | YAR015W | YBR182C | YAR015W | YAR018C |
| YBL023C | YBL023C | YBL034C | YAR018C | YBR199W | YAR018C | YBL042C |
| YBL034C | YBL034C | YBL042C | YBL023C | YBR202W | YAR018C | YBL106C |
| YBL042C | YBL042C | YBL085W | YBL034C | YBR297W | YBL023C | YBR002C |
| YBL085W | YBL085W | YBR021W | YBL042C | YCL051W | YBL042C | YBR038W |
| YBR021W | YBR021W | YBR038W | YBL085W | YCL064C | YBR021W | YBR039W |
| YBR038W | YBR038W | YBR055C | YBR021W | YCR088W | YBR021W | YBR046C |
| YBR055C | YBR055C | YBR085W | YBR038W | YDL101C | YBR021W | YBR085W |
| YBR150C | YBR085W | YBR150C | YBR055C | YDL120W | YBR026C | YBR095C |
| YBR160W | YBR150C | YBR160W | YBR085W | YDR063W | YBR038W | YBR102C |
| YBR177C | YBR160W | YBR177C | YBR150C | YDR196C | YBR085W | YBR150C |
| YBR182C | YBR177C | YBR182C | YBR160W | YDR393W | YBR085W | YBR172C |
| YBR202W | YBR182C | YBR202W | YBR182C | YDR456W | YBR092C | YBR182C |
| YBR210W | YBR202W | YBR210W | YBR202W | YEL072W | YBR150C | YBR182C |
| YBR271W | YBR210W | YBR271W | YBR210W | YER074W-A | YBR162W-A | YBR202W |
| YBR291C | YBR271W | YBR291C | YBR271W | YER146W | YBR182C | YBR210W |
| YBR296C | YBR291C | YBR296C | YBR291C | YER149C | YBR182C | YBR213W |
| YBR297W | YBR296C | YBR297W | YBR296C | YFR011C | YBR202W | YBR291C |
| YBR298C | YBR297W | YBR298C | YBR297W | YGL009C | YBR202W | YBR296C |
| YCL024W | YBR298C | YCL024W | YBR298C | YGL196W | YBR210W | YBR297W |
| YCR031C | YCL024W | YCR031C | YCL024W | YGR199W | YBR218C | YCL047C |
| YCR045C | YCR045C | YCR045C | YCR031C | YHR014W | YBR233W | YCL059C |
| YCR053W | YCR053W | YCR053W | YCR045C | YHR096C | YBR291C | YCR044C |
| YDL003W | YDL003W | YDL003W | YCR053W | YHR133C | YBR291C | YCR045C |
| YDL101C | YDL101C | YDL101C | YDL003W | YHR157W | YBR296C | YCR045C |
| YDL144C | YDL144C | YDL144C | YDL101C | YHR168W | YBR297W | YCR052W |
| YDL154W | YDL154W | YDL154W | YDL144C | YIL013C | YBR297W | YCR053W |
| YDL171C | YDL171C | YDL171C | YDL154W | YIL131C | YBR298C | YDL003W |
| YDL194W | YDL194W | YDL194W | YDL171C | YJL002C | YCL027W | YDL019C |
| YDR019C | YDR019C | YDR019C | YDL194W | YJR078W | YCL045C | YDL040C |
| YDR043C | YDR043C | YDR043C | YDR019C | YJR110W | YCR045C | YDL101C |
| YDR059C | YDR059C | YDR059C | YDR043C | YKL015W | YCR045C | YDL110C |
| YDR146C | YDR146C | YDR146C | YDR059C | YKL033W-A | YCR053W | YDL111C |
| YDR213W | YDR213W | YDR213W | YDR146C | YKL093W | YDL003W | YDL131W |
| YDR247W | YDR247W | YDR247W | YDR213W | YKL185W | YDL003W | YDL144C |
| YDR256C | YDR256C | YDR256C | YDR247W | YLR058C | YDL097C | YDL154W |
| YDR263C | YDR263C | YDR263C | YDR256C | YLR081W | YDL101C | YDR063W |
| YDR272W | YDR272W | YDR272W | YDR272W | YLR312C | YDL101C | YDR096W |
| YDR277C | YDR277C | YDR277C | YDR277C | YLR409C | YDL144C | YDR146C |
| YDR311W | YDR311W | YDR311W | YDR311W | YML042W | YDL154W | YDR169C |
| YDR317W | YDR317W | YDR317W | YDR317W | YMR018W | YDL154W | YDR175C |
| YDR374C | YDR374C | YDR374C | YDR374C | YMR189W | YDL182W | YDR213W |
| YDR375C | YDR375C | YDR375C | YDR375C | YNL041C | YDL226C | YDR225W |
| YDR393W | YDR393W | YDR393W | YDR393W | YNR015W | YDR007W | YDR229W |
| YDR403W | YDR403W | YDR403W | YDR403W | YOR036W | YDR043C | YDR256C |
| YDR451C | YDR451C | YDR451C | YDR451C | YOR066W | YDR142C | YDR263C |
| YDR507C | YDR507C | YDR507C | YDR507C | YOR229W | YDR146C | YDR272W |
| YDR524C | YDR524C | YDR524C | YDR524C | YPL011C | YDR146C | YDR277C |
| YDR528W | YDR528W | YDR528W | YDR528W | YPL088W | YDR204W | YDR294C |
| YDR532C | YDR532C | YDR532C | YDR532C | YPL104W | YDR238C | YDR295C |
| YEL017C-A | YEL017C-A | YEL017C-A | YEL017C-A | YPL264C | YDR247W | YDR311W |
| YEL030W | YEL030W | YEL030W | YEL030W |  | YDR256C | YDR317W |
| YEL039C | YEL039C | YEL039C | YEL039C |  | YDR263C | YDR317W |
| YEL042W | YEL042W | YEL042W | YEL042W |  | YDR263C | YDR341C |
| YEL072W | YEL072W | YEL072W | YEL072W |  | YDR272W | YDR347W |
| YER017C | YER017C | YER017C | YER017C |  | YDR277C | YDR349C |
| YER044C-A | YER044C-A | YER044C-A | YER044C-A |  | YDR306C | YDR374C |
| YER070W | YER070W | YER062C | YER070W |  | YDR311W | YDR389W |
| YER091C | YER091C | YER070W | YER091C |  | YDR317W | YDR393W |
| YER095W | YER095W | YER091C | YER095W |  | YDR357C | YDR395W |
| YER096W | YER096W | YER095W | YER096W |  | YDR358W | YDR403W |
| YER123W | YER123W | YER096W | YER123W |  | YDR374C | YDR408C |
| YER146W | YER146W | YER123W | YER146W |  | YDR375C | YDR424C |
| YER155C | YER155C | YER146W | YER155C |  | YDR393W | YDR451C |
| YFL052W | YFL052W | YER155C | YFL052W |  | YDR403W | YDR483W |
| YFL054C | YFL054C | YFL052W | YFL054C |  | YDR451C | YDR507C |
| YFR011C | YFR011C | YFL054C | YFR011C |  | YDR458C | YDR531W |
| YFR015C | YFR015C | YFR011C | YFR015C |  | YDR472W | YDR532C |
| YFR016C | YFR016C | YFR015C | YFR016C |  | YDR507C | YEL030W |
| YFR028C | YFR028C | YFR016C | YFR028C |  | YDR528W | YEL039C |
| YGL021W | YGL021W | YFR028C | YGL021W |  | YDR532C | YEL039C |
| YGL116W | YGL116W | YGL021W | YGL116W |  | YDR540C | YEL067C |
| YGL117W | YGL117W | YGL116W | YGL117W |  | YEL030W | YEL072W |
| YGL254W | YGL254W | YGL117W | YGL254W |  | YEL039C | YER009W |
| YGR087C | YGR087C | YGL254W | YGR087C |  | YEL039C | YER015W |
| YGR092W | YGR092W | YGR087C | YGR092W |  | YEL065W | YER044C-A |
| YGR129W | YGR129W | YGR092W | YGR129W |  | YEL072W | YER051W |
| YGR161C | YGR161C | YGR129W | YGR161C |  | YEL072W | YER068W |
| YGR177C | YGR177C | YGR161C | YGR177C |  | YER017C | YER070W |
| YGR199W | YGR199W | YGR177C | YGR199W |  | YER036C | YER091C |
| YGR221C | YGR221C | YGR199W | YGR221C |  | YER044C-A | YER095W |
| YGR240C | YGR240C | YGR221C | YGR240C |  | YER070W | YER096W |
| YGR248W | YGR248W | YGR240C | YGR248W |  | YER070W | YER126C |
| YGR281W | YGR281W | YGR248W | YGR281W |  | YER078C | YER128W |
| YGR289C | YGR289C | YGR281W | YGR289C |  | YER091C | YER140W |
| YHL026C | YHL026C | YGR289C | YHL026C |  | YER095W | YER146W |
| YHR014W | YHR014W | YHL026C | YHR014W |  | YER095W | YER151C |
| YHR032W | YHR032W | YHR014W | YHR032W |  | YER096W | YFL026W |
| YHR043C | YHR043C | YHR032W | YHR043C |  | YER096W | YFL047W |
| YHR048W | YHR048W | YHR043C | YHR048W |  | YER146W | YFL054C |
| YHR061C | YHR061C | YHR048W | YHR061C |  | YER146W | YFR011C |
| YHR096C | YHR096C | YHR061C | YHR096C |  | YER176W | YFR028C |
| YHR122W | YHR122W | YHR096C | YHR122W |  | YFL014W | YFR035C |
| YHR124W | YHR124W | YHR122W | YHR124W |  | YFL048C | YGL016W |
| YHR137W | YHR137W | YHR124W | YHR137W |  | YFL054C | YGL021W |
| YHR149C | YHR149C | YHR137W | YHR149C |  | YFL054C | YGL078C |
| YHR153C | YHR153C | YHR149C | YHR153C |  | YFR011C | YGL087C |
| YHR154W | YHR154W | YHR153C | YHR154W |  | YFR016C | YGL116W |
| YHR156C | YHR156C | YHR154W | YHR156C |  | YGL021W | YGL117W |
| YHR157W | YHR157W | YHR156C | YHR157W |  | YGL021W | YGL135W |
| YHR160C | YHR160C | YHR157W | YHR160C |  | YGL037C | YGL211W |
| YHR210C | YHR210C | YHR160C | YHR210C |  | YGL039W | YGL234W |
| YIL013C | YIL013C | YHR210C | YIL013C |  | YGL106W | YGL254W |
| YIL020C | YIL020C | YIL013C | YIL020C |  | YGL116W | YGR083C |
| YIL057C | YIL057C | YIL020C | YIL057C |  | YGL117W | YGR087C |
| YIL072W | YIL072W | YIL057C | YIL072W |  | YGL221C | YGR087C |
| YIL099W | YIL099W | YIL072W | YIL099W |  | YGL232W | YGR113W |
| YIL117C | YIL117C | YIL099W | YIL117C |  | YGL254W | YGR177C |
| YIL131C | YIL131C | YIL117C | YIL131C |  | YGR065C | YGR199W |
| YIL138C | YIL138C | YIL131C | YIL138C |  | YGR087C | YGR221C |
| YIL158W | YIL158W | YIL138C | YIL160C |  | YGR087C | YGR223C |
| YIL160C | YIL160C | YIL158W | YIR034C |  | YGR108W | YGR247W |
| YIR034C | YIR034C | YIL160C | YJL051W |  | YGR129W | YGR248W |
| YJL051W | YJL051W | YIR034C | YJL061W |  | YGR177C | YGR257C |
| YJL061W | YJL061W | YJL051W | YJL088W |  | YGR199W | YHR014W |
| YJL088W | YJL088W | YJL061W | YJL151C |  | YGR213C | YHR016C |
| YJL153C | YJL153C | YJL088W | YJL153C |  | YGR221C | YHR017W |
| YJL159W | YJL159W | YJL153C | YJL159W |  | YGR221C | YHR018C |
| YJL177W | YJL177W | YJL159W | YJL177W |  | YGR243W | YHR032W |
| YJL194W | YJL194W | YJL177W | YJL194W |  | YGR248W | YHR043C |
| YJL214W | YJL214W | YJL194W | YJL214W |  | YGR274C | YHR061C |
| YJL216C | YJL216C | YJL214W | YJL216C |  | YGR289C | YHR094C |
| YJR015W | YJR015W | YJL216C | YJR015W |  | YGR289C | YHR096C |
| YJR030C | YJR030C | YJR015W | YJR030C |  | YHR014W | YHR096C |
| YJR036C | YJR036C | YJR030C | YJR036C |  | YHR032W | YHR119W |
| YJR061W | YJR061W | YJR036C | YJR061W |  | YHR043C | YHR124W |
| YJR078W | YJR078W | YJR061W | YJR078W |  | YHR044C | YHR137W |
| YJR095W | YJR092W | YJR078W | YJR092W |  | YHR048W | YHR149C |
| YJR110W | YJR095W | YJR092W | YJR095W |  | YHR061C | YHR157W |
| YJR150C | YJR110W | YJR095W | YJR110W |  | YHR096C | YHR158C |
| YJR151C | YJR150C | YJR110W | YJR150C |  | YHR096C | YHR186C |
| YKL015W | YJR151C | YJR150C | YJR151C |  | YHR122W | YHR210C |
| YKL045W | YKL009W | YJR151C | YKL015W |  | YHR124W | YIL013C |
| YKL047W | YKL015W | YKL015W | YKL045W |  | YHR137W | YIL072W |
| YKL093W | YKL045W | YKL045W | YKL047W |  | YHR137W | YIL099W |
| YKL096W | YKL047W | YKL047W | YKL093W |  | YHR137W | YIL113W |
| YKL143W | YKL093W | YKL093W | YKL096W |  | YHR149C | YIL117C |
| YKL164C | YKL096W | YKL096W | YKL143W |  | YHR149C | YIL119C |
| YKL185W | YKL143W | YKL143W | YKL164C |  | YHR149C | YIL127C |
| YKL208W | YKL164C | YKL164C | YKL185W |  | YHR153C | YIL131C |
| YKR009C | YKL185W | YKL185W | YKL208W |  | YHR156C | YIL138C |
| YKR097W | YKL208W | YKL208W | YKR009C |  | YHR157W | YIL158W |
| YLL021W | YKR009C | YKR009C | YKR097W |  | YHR157W | YIL160C |
| YLL046C | YKR097W | YKR097W | YLL021W |  | YHR160C | YIR001C |
| YLL056C | YLL021W | YLL021W | YLL046C |  | YHR210C | YIR011C |
| YLR054C | YLL046C | YLL046C | YLL056C |  | YHR210C | YIR034C |
| YLR058C | YLL056C | YLL056C | YLR054C |  | YIL013C | YJL023C |
| YLR081W | YLR054C | YLR054C | YLR058C |  | YIL013C | YJL031C |
| YLR084C | YLR058C | YLR058C | YLR081W |  | YIL020C | YJL042W |
| YLR179C | YLR081W | YLR081W | YLR084C |  | YIL036W | YJL051W |
| YLR183C | YLR084C | YLR084C | YLR179C |  | YIL057C | YJL088W |
| YLR249W | YLR179C | YLR168C | YLR183C |  | YIL072W | YJL127C |
| YLR254C | YLR183C | YLR179C | YLR249W |  | YIL072W | YJL128C |
| YLR263W | YLR249W | YLR183C | YLR254C |  | YIL073C | YJL153C |
| YLR267W | YLR254C | YLR249W | YLR263W |  | YIL099W | YJL159W |
| YLR312C | YLR263W | YLR254C | YLR267W |  | YIL117C | YJL194W |
| YLR313C | YLR267W | YLR263W | YLR312C |  | YIL131C | YJL214W |
| YLR407W | YLR312C | YLR267W | YLR313C |  | YIL138C | YJL216C |
| YLR441C | YLR313C | YLR312C | YLR407W |  | YIL158W | YJR030C |
| YML042W | YLR407W | YLR313C | YLR441C |  | YIL158W | YJR030C |
| YML059C | YLR441C | YLR407W | YML042W |  | YIL158W | YJR061W |
| YML119W | YML042W | YLR441C | YML059C |  | YIL160C | YJR075W |
| YMR002W | YML059C | YML042W | YML119W |  | YIL166C | YJR078W |
| YMR006C | YML119W | YML059C | YMR002W |  | YIR034C | YJR082C |
| YMR018W | YMR002W | YML119W | YMR006C |  | YIR034C | YJR088C |
| YMR032W | YMR006C | YMR002W | YMR018W |  | YJL045W | YJR092W |
| YMR055C | YMR018W | YMR006C | YMR032W |  | YJL051W | YJR095W |
| YMR076C | YMR032W | YMR018W | YMR055C |  | YJL051W | YJR124C |
| YMR144W | YMR055C | YMR032W | YMR076C |  | YJL088W | YJR150C |
| YMR189W | YMR076C | YMR055C | YMR144W |  | YJL088W | YJR151C |
| YMR240C | YMR144W | YMR076C | YMR189W |  | YJL093C | YKL003C |
| YMR278W | YMR189W | YMR144W | YMR240C |  | YJL145W | YKL015W |
| YMR280C | YMR240C | YMR177W | YMR266W |  | YJL153C | YKL047W |
| YNL012W | YMR278W | YMR189W | YMR278W |  | YJL159W | YKL052C |
| YNL014W | YMR280C | YMR240C | YMR280C |  | YJL159W | YKL093W |
| YNL037C | YNL012W | YMR278W | YNL012W |  | YJL162C | YKL096W |
| YNL058C | YNL014W | YMR280C | YNL014W |  | YJL194W | YKL129C |
| YNL112W | YNL037C | YNL012W | YNL037C |  | YJL194W | YKL130C |
| YNL183C | YNL041C | YNL014W | YNL041C |  | YJL214W | YKL143W |
| YNL196C | YNL058C | YNL037C | YNL058C |  | YJL216C | YKL173W |
| YNL208W | YNL112W | YNL041C | YNL112W |  | YJL216C | YKL185W |
| YNL234W | YNL183C | YNL058C | YNL183C |  | YJR036C | YKL186C |
| YNL270C | YNL196C | YNL112W | YNL196C |  | YJR036C | YKL208W |
| YNL279W | YNL208W | YNL183C | YNL208W |  | YJR036C | YKL218C |
| YNL306W | YNL234W | YNL196C | YNL234W |  | YJR039W | YKR005C |
| YNR002C | YNL259C | YNL208W | YNL270C |  | YJR061W | YKR009C |
| YNR015W | YNL270C | YNL234W | YNL306W |  | YJR078W | YKR017C |
| YNR061C | YNL279W | YNL270C | YNR002C |  | YJR092W | YKR037C |
| YOL007C | YNL306W | YNL279W | YNR015W |  | YJR095W | YKR051W |
| YOL014W | YNR002C | YNL306W | YNR061C |  | YJR104C | YKR084C |
| YOL017W | YNR015W | YNR002C | YOL007C |  | YJR123W | YKR097W |
| YOL066C | YNR061C | YNR015W | YOL014W |  | YJR150C | YLL003W |
| YOL084W | YOL007C | YNR061C | YOL017W |  | YJR151C | YLL021W |
| YOL091W | YOL014W | YOL007C | YOL034W |  | YJR151C | YLL048C |
| YOL104C | YOL017W | YOL014W | YOL066C |  | YKL015W | YLL056C |
| YOL151W | YOL034W | YOL017W | YOL084W |  | YKL015W | YLR018C |
| YOR009W | YOL066C | YOL030W | YOL091W |  | YKL047W | YLR028C |
| YOR019W | YOL084W | YOL034W | YOL104C |  | YKL093W | YLR054C |
| YOR058C | YOL091W | YOL066C | YOL151W |  | YKL093W | YLR067C |
| YOR062C | YOL104C | YOL084W | YOR009W |  | YKL096W | YLR081W |
| YOR066W | YOL151W | YOL091W | YOR019W |  | YKL096W | YLR084C |
| YOR083W | YOR009W | YOL100W | YOR058C |  | YKL143W | YLR100W |
| YOR100C | YOR019W | YOL104C | YOR062C |  | YKL185W | YLR106C |
| YOR167C | YOR058C | YOL151W | YOR066W |  | YKL186C | YLR121C |
| YOR180C | YOR062C | YOR009W | YOR083W |  | YKL208W | YLR178C |
| YOR185C | YOR066W | YOR019W | YOR100C |  | YKL208W | YLR183C |
| YOR229W | YOR083W | YOR058C | YOR167C |  | YKR009C | YLR238W |
| YOR287C | YOR100C | YOR062C | YOR180C |  | YKR009C | YLR249W |
| YOR291W | YOR167C | YOR066W | YOR185C |  | YKR009C | YLR254C |
| YPL066W | YOR180C | YOR083W | YOR229W |  | YKR027W | YLR263W |
| YPL090C | YOR185C | YOR100C | YOR287C |  | YKR097W | YLR292C |
| YPL104W | YOR229W | YOR167C | YOR291W |  | YLL021W | YLR305C |
| YPL112C | YOR287C | YOR180C | YPL066W |  | YLL040C | YLR312C |
| YPL133C | YOR291W | YOR185C | YPL090C |  | YLL046C | YLR409C |
| YPL189W | YPL066W | YOR229W | YPL104W |  | YLL056C | YLR447C |
| YPL200W | YPL090C | YOR287C | YPL112C |  | YLL056C | YML004C |
| YPL201C | YPL104W | YOR291W | YPL133C |  | YLR054C | YML031W |
| YPL206C | YPL112C | YPL066W | YPL189W |  | YLR054C | YML042W |
| YPL212C | YPL133C | YPL090C | YPL200W |  | YLR058C | YML059C |
| YPL252C | YPL163C | YPL104W | YPL201C |  | YLR058C | YML120C |
| YPL274W | YPL189W | YPL112C | YPL206C |  | YLR066W | YMR006C |
| YPR006C | YPL200W | YPL133C | YPL212C |  | YLR081W | YMR018W |
| YPR013C | YPL201C | YPL189W | YPL242C |  | YLR084C | YMR032W |
| YPR015C | YPL206C | YPL200W | YPL252C |  | YLR183C | YMR047C |
| YPR026W | YPL212C | YPL201C | YPL274W |  | YLR183C | YMR055C |
| YPR109W | YPL242C | YPL206C | YPR006C |  | YLR228C | YMR076C |
| YPR119W | YPL252C | YPL212C | YPR013C |  | YLR249W | YMR091C |
|  | YPL274W | YPL242C | YPR015C |  | YLR254C | YMR114C |
|  | YPR006C | YPL252C | YPR026W |  | YLR263W | YMR182C |
|  | YPR013C | YPL274W | YPR109W |  | YLR263W | YMR187C |
|  | YPR015C | YPR006C | YPR119W |  | YLR267W | YMR189W |
|  | YPR026W | YPR013C |  |  | YLR297W | YMR207C |
|  | YPR109W | YPR015C |  |  | YLR312C | YMR214W |
|  | YPR119W | YPR026W |  |  | YLR313C | YMR261C |
|  |  | YPR109W |  |  | YLR407W | YMR270C |
|  |  | YPR119W |  |  | YLR440C | YMR311C |
|  |  |  |  |  | YLR443W | YNL005C |
|  |  |  |  |  | YML004C | YNL014W |
|  |  |  |  |  | YML028W | YNL041C |
|  |  |  |  |  | YML042W | YNL058C |
|  |  |  |  |  | YML042W | YNL085W |
|  |  |  |  |  | YML059C | YNL108C |
|  |  |  |  |  | YMR006C | YNL152W |
|  |  |  |  |  | YMR018W | YNL176C |
|  |  |  |  |  | YMR018W | YNL196C |
|  |  |  |  |  | YMR032W | YNL208W |
|  |  |  |  |  | YMR047C | YNL234W |
|  |  |  |  |  | YMR055C | YNL270C |
|  |  |  |  |  | YMR076C | YNL306W |
|  |  |  |  |  | YMR100W | YNR015W |
|  |  |  |  |  | YMR153W | YNR044W |
|  |  |  |  |  | YMR188C | YOL007C |
|  |  |  |  |  | YMR189W | YOL017W |
|  |  |  |  |  | YMR189W | YOL034W |
|  |  |  |  |  | YMR280C | YOL066C |
|  |  |  |  |  | YMR290C | YOL084W |
|  |  |  |  |  | YNL014W | YOL091W |
|  |  |  |  |  | YNL014W | YOL104C |
|  |  |  |  |  | YNL024C | YOL151W |
|  |  |  |  |  | YNL041C | YOR006C |
|  |  |  |  |  | YNL053W | YOR009W |
|  |  |  |  |  | YNL058C | YOR019W |
|  |  |  |  |  | YNL117W | YOR058C |
|  |  |  |  |  | YNL175C | YOR066W |
|  |  |  |  |  | YNL192W | YOR083W |
|  |  |  |  |  | YNL196C | YOR099W |
|  |  |  |  |  | YNL196C | YOR100C |
|  |  |  |  |  | YNL208W | YOR180C |
|  |  |  |  |  | YNL234W | YOR185C |
|  |  |  |  |  | YNL234W | YOR259C |
|  |  |  |  |  | YNL261W | YOR287C |
|  |  |  |  |  | YNL265C | YOR301W |
|  |  |  |  |  | YNL270C | YPL007C |
|  |  |  |  |  | YNL306W | YPL066W |
|  |  |  |  |  | YNL306W | YPL090C |
|  |  |  |  |  | YNR002C | YPL100W |
|  |  |  |  |  | YNR002C | YPL104W |
|  |  |  |  |  | YNR015W | YPL112C |
|  |  |  |  |  | YNR015W | YPL121C |
|  |  |  |  |  | YOL007C | YPL183C |
|  |  |  |  |  | YOL017W | YPL200W |
|  |  |  |  |  | YOL017W | YPL201C |
|  |  |  |  |  | YOL038W | YPL203W |
|  |  |  |  |  | YOL066C | YPL235W |
|  |  |  |  |  | YOL084W | YPL252C |
|  |  |  |  |  | YOL088C | YPR006C |
|  |  |  |  |  | YOL091W | YPR015C |
|  |  |  |  |  | YOL091W | YPR026W |
|  |  |  |  |  | YOL104C | YPR119W |
|  |  |  |  |  | YOL104C | YPR125W |
|  |  |  |  |  | YOL154W |  |
|  |  |  |  |  | YOR009W |  |
|  |  |  |  |  | YOR009W |  |
|  |  |  |  |  | YOR016C |  |
|  |  |  |  |  | YOR019W |  |
|  |  |  |  |  | YOR058C |  |
|  |  |  |  |  | YOR058C |  |
|  |  |  |  |  | YOR066W |  |
|  |  |  |  |  | YOR077W |  |
|  |  |  |  |  | YOR083W |  |
|  |  |  |  |  | YOR100C |  |
|  |  |  |  |  | YOR100C |  |
|  |  |  |  |  | YOR100C |  |
|  |  |  |  |  | YOR167C |  |
|  |  |  |  |  | YOR180C |  |
|  |  |  |  |  | YOR185C |  |
|  |  |  |  |  | YOR185C |  |
|  |  |  |  |  | YOR210W |  |
|  |  |  |  |  | YOR260W |  |
|  |  |  |  |  | YOR287C |  |
|  |  |  |  |  | YOR291W |  |
|  |  |  |  |  | YOR316C |  |
|  |  |  |  |  | YOR384W |  |
|  |  |  |  |  | YPL006W |  |
|  |  |  |  |  | YPL065W |  |
|  |  |  |  |  | YPL066W |  |
|  |  |  |  |  | YPL090C |  |
|  |  |  |  |  | YPL104W |  |
|  |  |  |  |  | YPL104W |  |
|  |  |  |  |  | YPL112C |  |
|  |  |  |  |  | YPL112C |  |
|  |  |  |  |  | YPL112C |  |
|  |  |  |  |  | YPL171C |  |
|  |  |  |  |  | YPL179W |  |
|  |  |  |  |  | YPL200W |  |
|  |  |  |  |  | YPL200W |  |
|  |  |  |  |  | YPL201C |  |
|  |  |  |  |  | YPL201C |  |
|  |  |  |  |  | YPL212C |  |
|  |  |  |  |  | YPL252C |  |
|  |  |  |  |  | YPL274W |  |
|  |  |  |  |  | YPR006C |  |
|  |  |  |  |  | YPR013C |  |
|  |  |  |  |  | YPR015C |  |
|  |  |  |  |  | YPR022C |  |
|  |  |  |  |  | YPR026W |  |
|  |  |  |  |  | YPR089W |  |
|  |  |  |  |  | YPR109W |  |
|  |  |  |  |  | YPR109W |  |
|  |  |  |  |  | YPR119W |  |

**Table S3:**
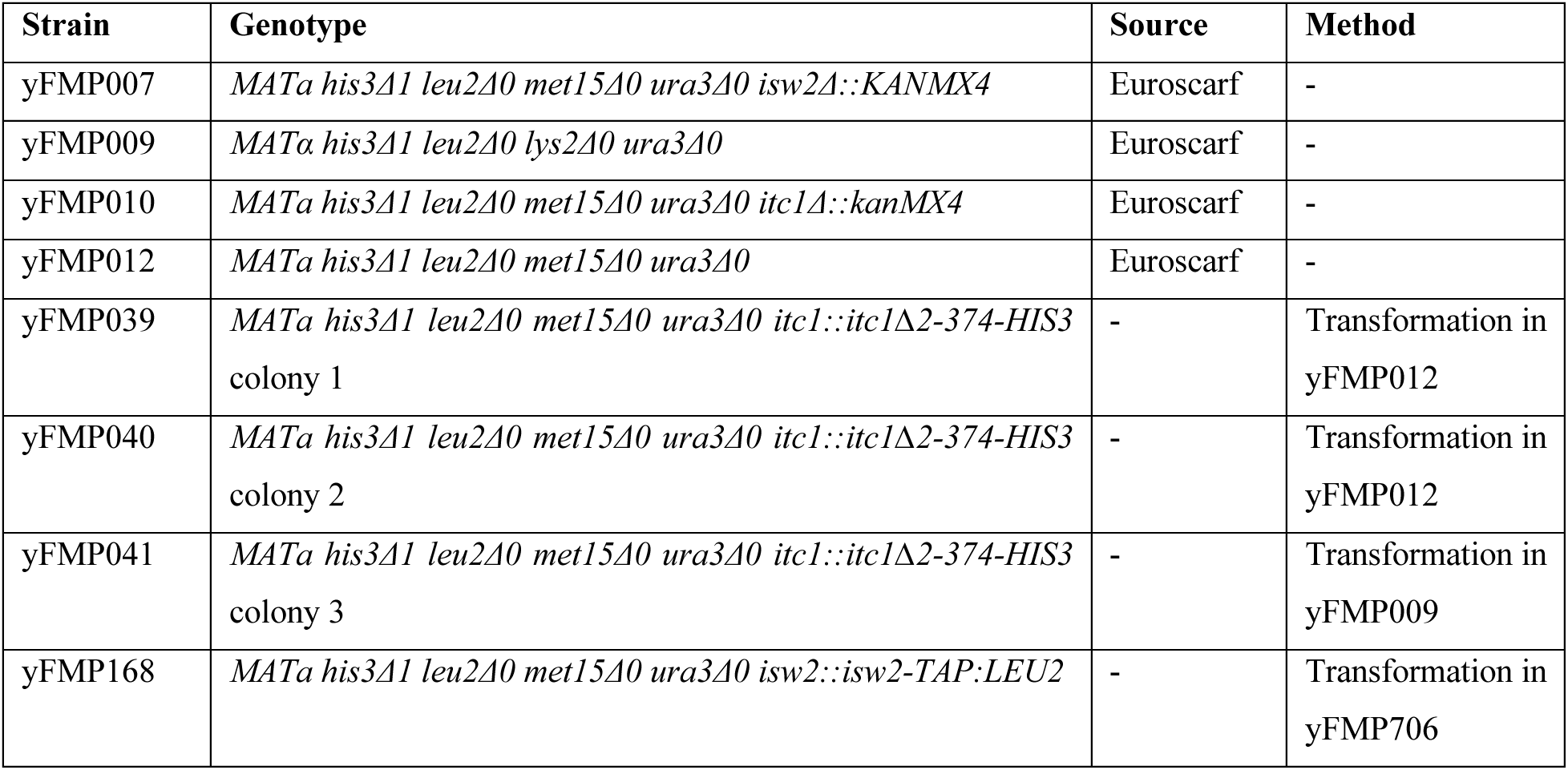
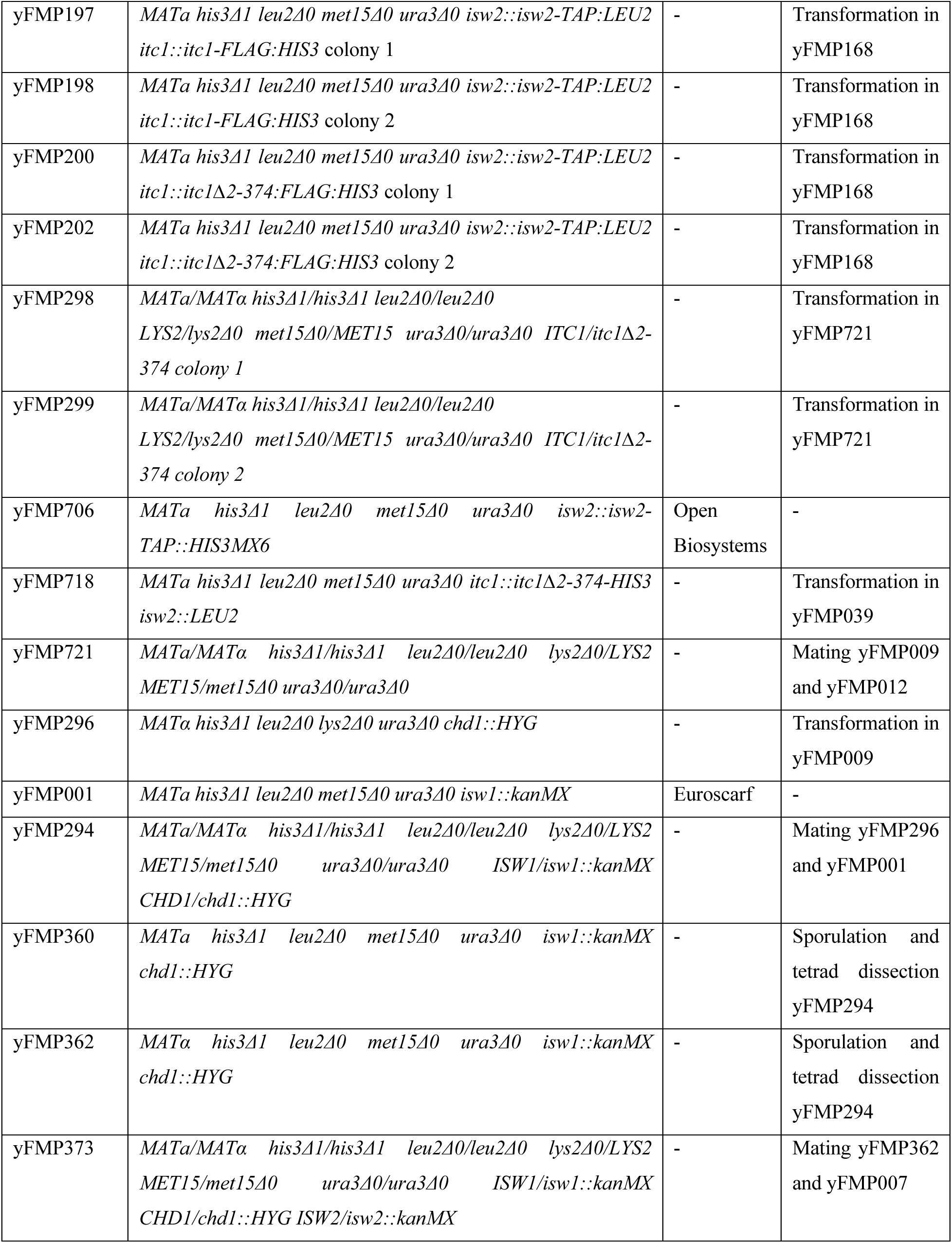
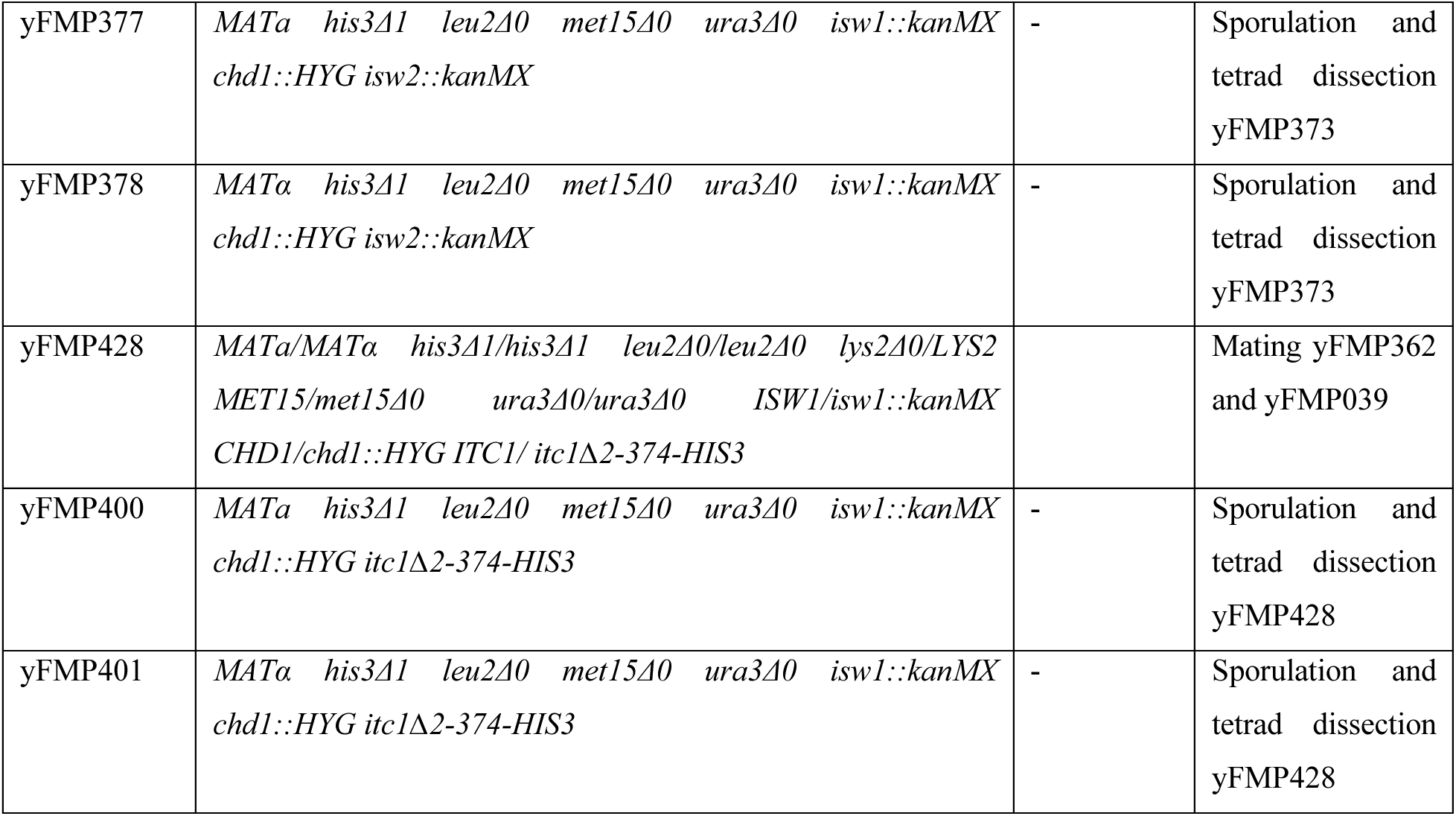
List of yeast strains.

**Table S4:**
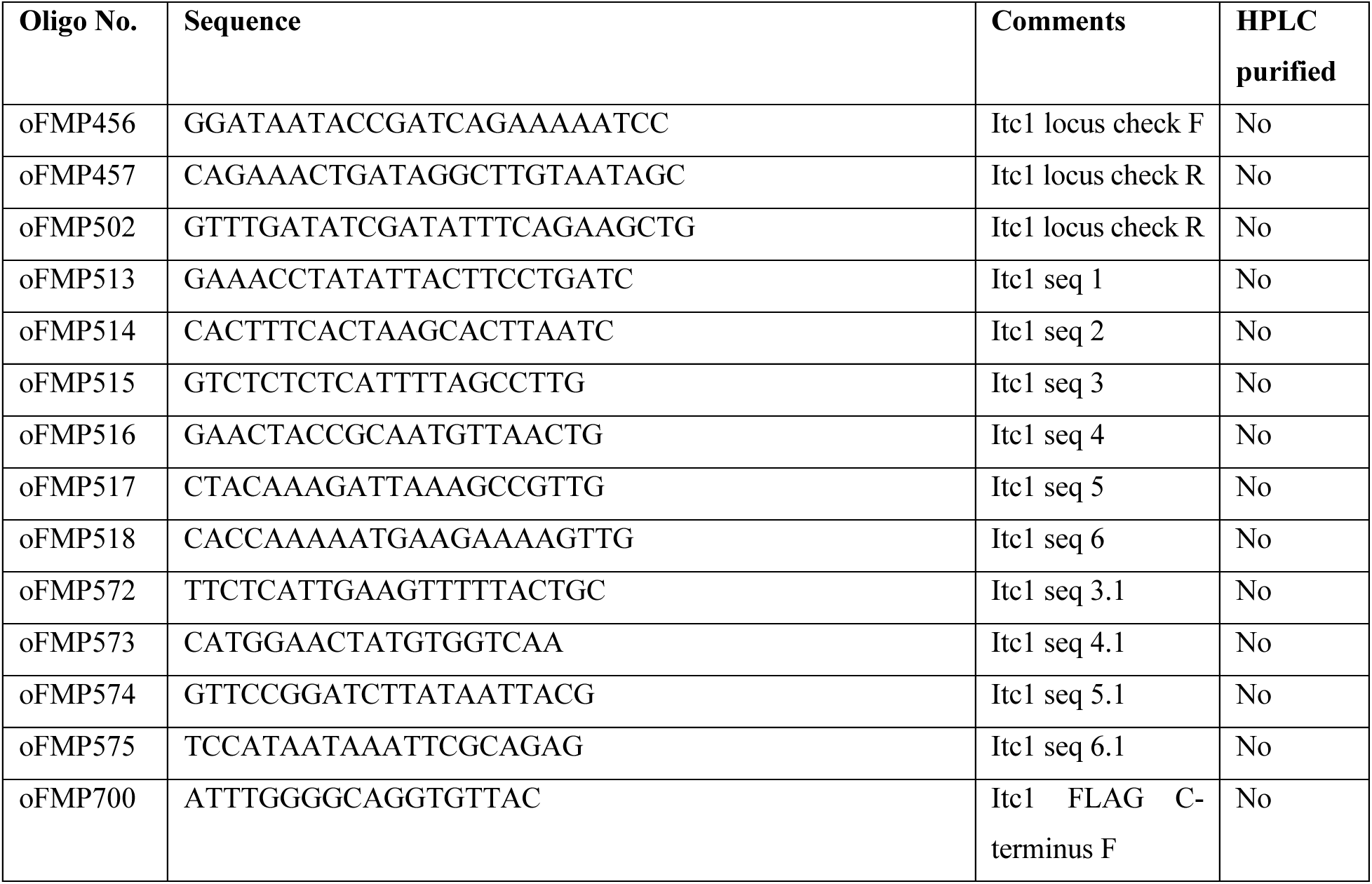
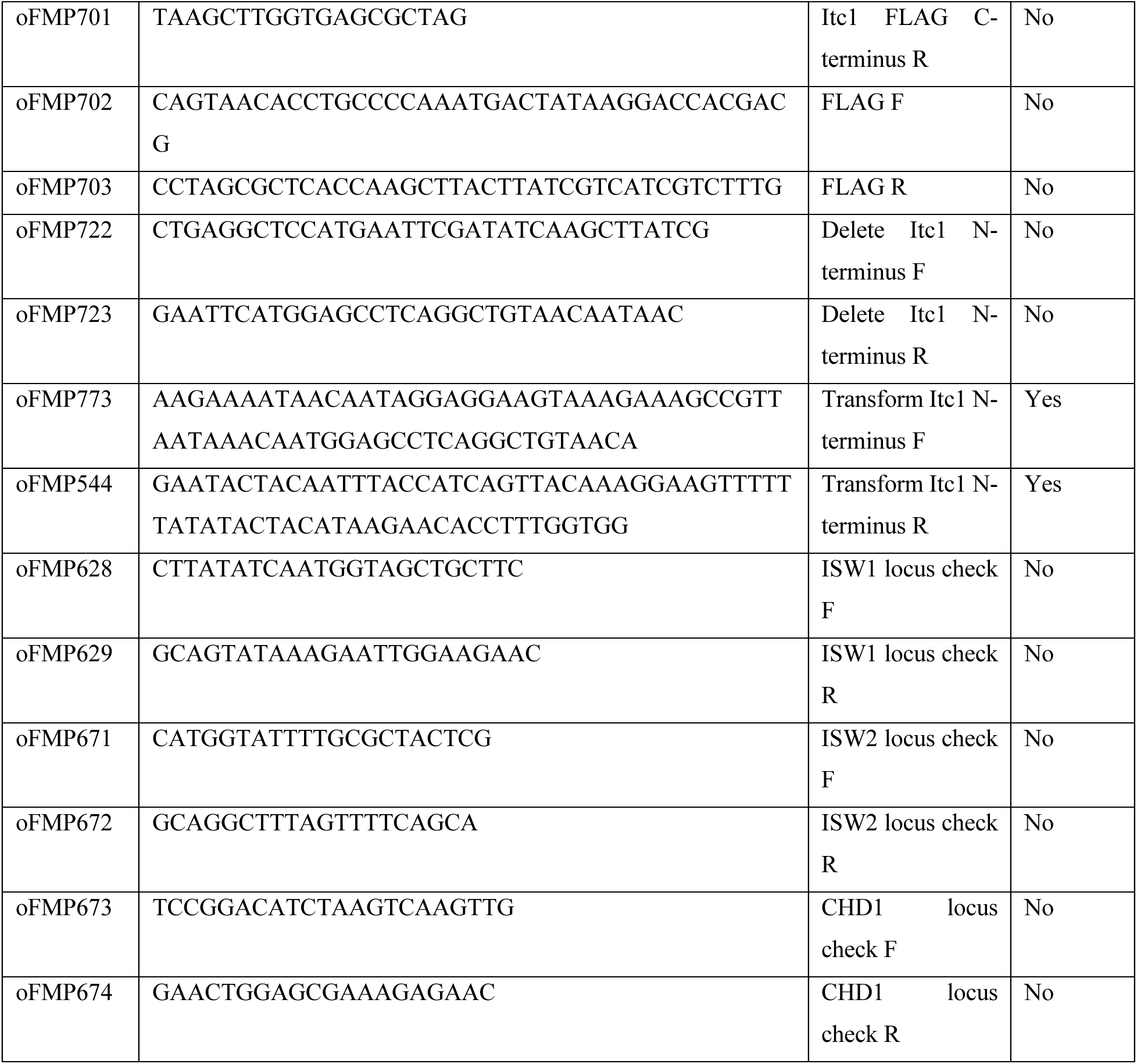
List of oligonucleotides.

